# Early life high fructose exposure disrupts microglia function and impedes neurodevelopment

**DOI:** 10.1101/2023.08.14.553242

**Authors:** Zhaoquan Wang, Allie Lipshutz, Zong-Lin Liu, Alissa J. Trzeciak, Isabella C. Miranda, Celia Martínez de la Torre, Tanya Schild, Tomi Lazarov, Waleska Saitz Rojas, Pedro H. V. Saavedra, Jesús E. Romero-Pichardo, Ann Baako, Frederic Geissmann, Giuseppe Faraco, Li Gan, Jon Iker Etchegaray, Christopher D. Lucas, Christopher N. Parkhurst, Melody Y. Zeng, Kayvan R. Keshari, Justin S. A. Perry

**Affiliations:** Immunology Program, Sloan Kettering Institute, Memorial Sloan Kettering Cancer Center, New York, NY, USA; Immunology and Microbial Pathogenesis Program, Weill Cornell Medicine, New York, NY, USA; Division of Pulmonary and Critical Care Medicine, Weill Cornell Medicine, New York, NY, USA; Department of Radiology, Sloan Kettering Institute, Memorial Sloan Kettering Cancer Center, New York, NY, USA; Molecular Pharmacology Program, Sloan Kettering Institute, Memorial Sloan Kettering Cancer Center, New York, NY, USA; Louis V. Gerstner Jr. Graduate School of Biomedical Sciences, Memorial Sloan Kettering Cancer Center, New York, NY, USA; Neuroscience Graduate Program, Weill Cornell Medicine, New York, NY, USA; Feil Family Brain and Mind Research Institute, Weill Cornell Medicine, New York, NY, USA; Helen and Robert Appel Alzheimer’s Disease Research Institute, Feil Family Brain and Mind Research Institute, Weill Cornell Medicine, New York, NY, USA; Department of Pathology and Immunology, Washington University School of Medicine, St. Louis, MO, USA; University of Edinburgh Centre for Inflammation Research, Queen’s Medical Research Institute, Edinburgh BioQuarter, Edinburgh, Scotland, UK; Institute for Regeneration and Repair, Edinburgh BioQuarter, Edinburgh, Scotland, UK; Gale and Ira Drukier Institute for Children’s Health, Weill Cornell Medicine, New York, NY, USA; Department of Pediatrics, Neonatal Medicine, Weill Cornell Medicine, New York, NY, USA

## Abstract

Despite the success of fructose as a low-cost food additive, recent epidemiological evidence suggests that high fructose consumption by pregnant mothers or during adolescence is associated with disrupted neurodevelopment^1–7^. An essential step in appropriate mammalian neurodevelopment is the synaptic pruning and elimination of newly-formed neurons by microglia, the central nervous system’s (CNS) resident professional phagocyte^8–10^. Whether early life high fructose consumption affects microglia function and if this directly impacts neurodevelopment remains unknown. Here, we show that both offspring born to dams fed a high fructose diet and neonates exposed to high fructose exhibit decreased microglial density, increased uncleared apoptotic cells, and decreased synaptic pruning *in vivo*. Importantly, deletion of the high affinity fructose transporter SLC2A5 (GLUT5) in neonates completely reversed microglia dysfunction, suggesting that high fructose directly affects neonatal development. Mechanistically, we found that high fructose treatment of both mouse and human microglia suppresses synaptic pruning and phagocytosis capacity which is fully reversed in GLUT5-deficient microglia. Using a combination of *in vivo* and *in vitro* nuclear magnetic resonance- and mass spectrometry-based fructose tracing, we found that high fructose drives significant GLUT5-dependent fructose uptake and catabolism, rewiring microglia metabolism towards a hypo-phagocytic state. Importantly, mice exposed to high fructose as neonates exhibited cognitive defects and developed anxiety-like behavior which were rescued in GLUT5-deficient animals. Our findings provide a mechanistic explanation for the epidemiological observation that early life high fructose exposure is associated with increased prevalence of adolescent anxiety disorders.

## Introduction

High fructose corn syrup (HFCS)-containing foods and beverages have faced increasing scrutiny over the past few decades, as the health hazards of overconsumption continue to be highlighted^11,12^. Several studies implicate high fructose as a key metabolic disease risk factor, contributing to the development of obesity, diabetes, and cardiovascular disease^13–15^. Recently, multiple epidemiological studies have suggested that high fructose consumption by pregnant mothers or during adolescence negatively impacts neurodevelopment, including increased risk of mood and anxiety disorder development^1–7,16–20^. Experiments in rodents have shown that chronic high fructose consumption can contribute to the development of cardiometabolic disease, specifically involving small intestine and liver fructose catabolism (fructolysis)^14,21–25^. On the other hand, a series of independent studies found that fructolysis can occur in the central nervous system (CNS)^26–29^. Whether the neurodevelopmental effects observed in epidemiological studies are a result of fructose metabolism in the CNS and how high fructose affects CNS cellular function remains unknown.

In mammals, neurodevelopment occurs through well-defined stages that are tightly temporally controlled^30,31^. In particular, neurons proliferate and form synapses which, if unreinforced, must be eliminated through a process termed synaptic pruning^8,9^. Successful synaptic pruning is accompanied by programmed cell death of neurons^32–35^. Both synaptic pruning and the phagocytic clearance of dying neurons (termed efferocytosis) are executed by microglia, the CNS’s resident macrophage and professional phagocyte^10,36–42^. Importantly, disrupted synaptic pruning or the removal of dying neurons can have catastrophic consequences on neurodevelopment in animals, including the development of anxiety-like behavior^43–48^. Here, we investigated the hypothesis that early life high fructose directly affects essential microglia phagocytic processes and whether early life high fructose causes defects in learning and the development of anxiety-like behavior.

## Results

### Early life high fructose exposure suppresses microglial phagocytosis *in vivo*

High fructose consumption, either by pregnant mothers or during adolescence, is correlated with neurodevelopmental sequalae including mood and anxiety disorder development. To test whether high fructose exposure directly affects microglia, the CNS-resident phagocyte essential for neurodevelopment, we delivered fructose to neonates using intragastric injection (Extended Data Fig. 1a). Strikingly, we observed significant reductions in microglia numbers and increased numbers of uncleared dying cells in neonates treated with high fructose compared to vehicle-treated neonates (Extended Data Fig. 1b-d).

We next queried whether this phenotype would extend to neonates born to dams on high fructose. This is especially important, because a recent study found that fructose is transmitted via breast milk and that increased maternal fructose consumption (e.g., via sugar-sweetened beverages or juice) correlates with negative neurodevelopmental outcomes^4,49^. Previous studies of high fructose used diets containing supraphysiological levels of fructose (up to 60% of total kcal)^50–52^. Instead, we placed mouse dams on isocaloric diets containing either 0 kcal% sucrose/0 kcal% fructose (control diet, CD) or clinically-relevant 15 kcal% high fructose diet (HF) that more accurately reflects average consumption in the United States^53^ (**Fig. 1A**). We then analyzed the prefrontal cortex (PFC) in brains of offspring at perinatal day 7 (P7), which represents the key period where microglia-mediated synaptic pruning and dying neuron clearance peaks^35^. We found that neonates born to and nursed by dams on HF diet had significantly fewer microglia compared to mice born to and nursed by dams given control diet (**Fig. 1b, c**). We also observed altered cell morphology in neonates born to and nursed by dams on HF diet. To objectively quantify this, we developed a battery of approaches based on literature defining microglia morphological changes indicative of cellular function (and dysfunction; Extended Data Fig. 2a)^54–61^. Using this battery, we found that microglia in neonates born to and nursed by dams on HF diet exhibit decreased soma size, decreased major axis length, and increased soma roundness (**Fig. 1d**), all previously shown as phenotypes associated with quiescent, non-phagocytic microglia^10,62,63^. Contrarily, we did not detect significant changes in microglia branching or branch junctions, which are not associated with neonatal microglia phagocytic activity (Extended Data Fig. 2b).

**Fig. 1:**
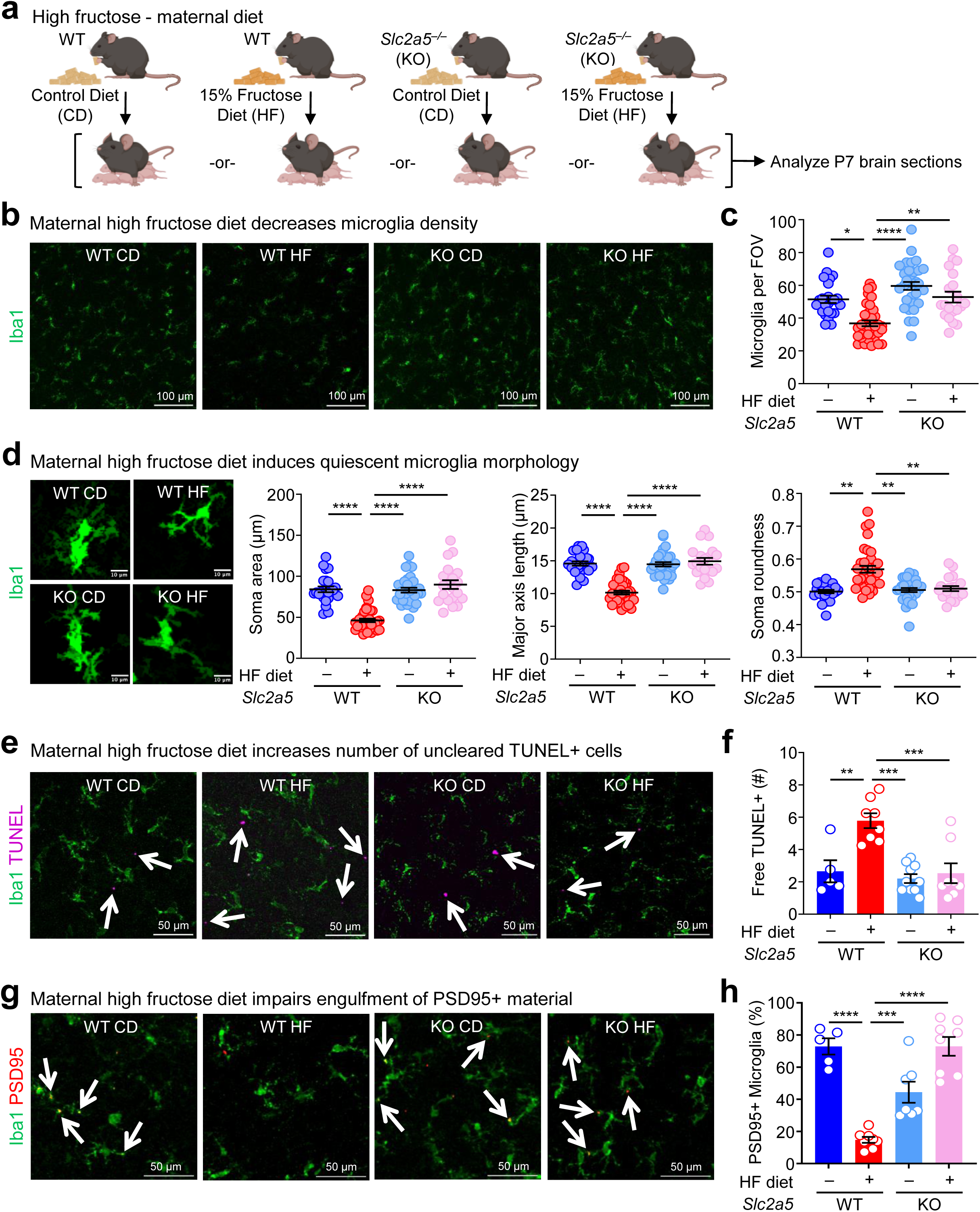
Early life high fructose exposure suppresses microglial phagocytosis *in vivo*. **(a)** <u>Early life dietary high fructose exposure in WT and *Slc2a5^−/−^* mice.</u> Schematic of wildtype (WT) and *Slc2a5^−/−^* (KO) dams placed on 0 kcal% sucrose control diet (CD) or 15 kcal% fructose diet (HF) for at least one week prior to pregnancy and maintained through gestation and lactation. Brains from postnatal day 7 (P7) mice were collected and analyzed using confocal microscopy. **(b, c)** <u>Maternal high fructose diet decreases microglia density.</u> Representative images **(b)** and quantitation **(c)** of microglia (Iba1+) numbers in the prefrontal cortex from WT and KO P7 mice exposed to CD or HF. Number of microglia were quantified using Iba1 staining, with four fields of view (FOVs) per animal plotted. Images are representative of six WT mice on CD (blue), nine WT mice on HF (red), eight KO mice on CD (light blue), and five KO mice on HF (pink). Scale bar, 100 µm. **(d)** <u>Maternal high fructose diet induces quiescent microglia morphology</u>. (Left) Representative images of microglia in the prefrontal cortex from WT and KO P7 mice exposed to maternal CD or HF as detailed in **(a)** used to analyze microglia morphology (see Extended Data Fig. 2a) from experiments in **(b)**. Scale bar, 10 µm. (Right) Quantitation of soma area (left), soma major axis length (middle), and soma roundness (right) from microglial morphological analysis of four FOVs per animal. Each dot represents an individual FOV with all cells in each FOV contributing to the plotted average. Scale bar, 10 µm. **(e, f)** <u>Maternal high fructose diet increases number of uncleared TUNEL+ cells.</u> Representative images **(e)** and quantitation **(f)** of uncleared (‘free’) TUNEL+ puncta in the prefrontal cortex from WT and KO P7 mice exposed to CD or HF in **(b)**. Free and bound/internalized TUNEL+ puncta (see Extended Data Fig. 3c) were identified in four FOVs per mouse for each condition and the average of the FOVs for each mouse is shown. Images shown are from zoomed in sections of 424.27 µm^2^ FOV, consistent with image sizes in **(b)**. Scale bar, 50 µm. **(g, h)** <u>Maternal high fructose diet impairs engulfment of PSD95+ material.</u> Representative images **(g)** and quantitation **(h)** of microglia phagocytosis of PSD95+ material in the prefrontal cortex from WT and KO P7 mice exposed to maternal CD or HF in **(b)**. The number of microglia with bound/internalized PSD95+ material was tabulated from four FOVs per mouse for each condition and the average of the FOVs for each mouse is shown. Images shown are from zoomed in sections of 424.27 µm^2^ FOV, consistent with image sizes in **(b)**. Scale bar, 50 µm. Data are shown as mean ± SEM. Significance was determined using two-way ANOVA. \**p* < .05, \*\**p* < .01, \*\*\**p* < 0.001, \*\*\*\**p* < .0001. Statistical analyses were performed using the averages calculated from the FOVs for each animal, not pseudo-replications.

Given the significant decrease in microglia numbers and altered cell morphology, we hypothesized that neonates born to and nursed by dams on HF diet would also display evidence of phagocytosis defects. Indeed, we observed a significant increase in uncleared (‘free’) pyknotic (dying cell) nuclei in neonates born to and nursed by dams on HF diet (Extended Data Fig. 3a, b). To more accurately determine if microglia phagocytosis is disrupted in neonates born to and nursed by dams on HF diet, we took two complimentary approaches. First, we used the terminal deoxynucleotidyl transferase dUTP nick end labeling (TUNEL) assay to identify uncleared dead cells^64,65^ (Extended Data Fig. 3c). Second, we analyzed microglia for the presence of postsynaptic density protein 95 (PSD95) material, an established readout for synaptic pruning^8^. Similar to our results showing a significant increase in uncleared pyknotic nuclei, neonates born to and nursed by dams on HF diet exhibited a significant increase in uncleared TUNEL-positive cells (**Fig. 1e, f**). Additionally, significantly fewer microglia in neonates born to and nursed by dams on HF diet contained PSD95 material and contained significantly less engulfed PSD95+ material on a per-cell basis (**Fig. 1g, h** and Extended Data Fig. 3d), further supporting the hypothesis that HF diet suppresses essential microglia phagocytic functions. Taken together, our results suggest that neonates exposed to high fructose have disrupted microglia form and function, including a significant reduction in essential phagocytic activities, *in vivo*.

### The effect of early life high fructose exposure depends on GLUT5-mediated fructose transport in neonates

Fructose transport is achieved by the facilitative glucose transport family members GLUT2 (*Slc2a2*) and GLUT5 (*Slc2a5*)^25^. In adults, GLUT5 mediates transport of dietary fructose from the intestinal lumen across the brush border membrane, followed by GLUT2-mediated transport across the basolateral membrane into the blood^24^. Contrarily, gestational and pre-weaning neonates express little if any GLUT5 but stably express GLUT2, including reportedly in the brush-border membrane^66–71^, suggesting that fructose delivered to pre-weaning neonates could gain access to the brain independent of GLUT5 in the intestinal lumen. Interestingly, GLUT5 was previously identified as a core microglia signature gene/protein together with TMEM119 and P2Y12^72^. Analysis of previously published informatics data revealed that microglia (including neonatal microglia) are the only immune cell in mice and humans that express GLUT5^73,74^ (Extended Data Fig. 3e) and are the only CNS-resident cell in mice and humans to express GLUT5^75–78^ (Extended Data Fig. 3f). Given this series of observations, we sought to determine if neonatal GLUT5 is necessary for the microglia phenotypes observed in high fructose-treated mice. To this end, we used *Slc2a5* deficient mice in which exons 1-4 are deleted^79^ (*Slc2a5*^−/−^ mice; Extended Data Fig. 4a, b) to performed high fructose diet experiments (**Fig. 1a**). Strikingly, microglia numbers, morphology, and phagocytic capacity were completely reversed in neonates lacking GLUT5 born to and nursed by dams on HF diet compared to wildtype neonates born to and nursed by dams on HF diet (**Fig. 1b-h**). Thus, neonatal GLUT5 facilitates the negative effects of early life high fructose on the developing CNS.

### High fructose exposure directly suppresses microglia phagocytosis

Given our finding that high fructose affects neonatal microglia form and function *in vivo* and microglia singularly express SLC2A5 (GLUT5) in both the CNS and the immune system at large, we speculated that high fructose was acting directly on microglia via GLUT5. In support of this hypothesis, microglia from neonates born to and nursed by dams on HF diet exhibited a significant increase in *Slc2a5* (mRNA, **Fig. 2a**) and SLC2A5 (protein, **Fig. 2b**) expression compared to microglia from neonates born to and nursed by dams on control diet. To test this hypothesis directly, we next tested the effect of physiologically-relevant high fructose on primary microglia function at a physiological level of oxygen *in vitro*. Specifically, we used a dedicated chamber system to culture microglia at a constant 3.5% O_2_^80–82^, consistent with the oxygen level observed in the CNS parenchyma where microglia reside (∼3-4% O_2_)^83–85^. Furthermore, we used an established media formulation that excludes serum because serum exposure has been shown to artificially affect microglia form and function (especially phagocytosis)^86^. Primary microglia isolated from P2-P4 neonates were cultured in low (1 mM) and high (5 mM) fructose with a hexose osmolarity control (mannitol) in fixed physiological glucose (5 mM) for 7 days (Extended Data Fig. 5a). Similar to our *in vivo* observations, mixed glial cultures cultured with high fructose generated fewer numbers of primary mouse microglia (Extended Data Fig. 5b). We next assessed microglia phagocytosis of two key targets: synaptic terminals (synaptosomes) and apoptotic neurons (Extended Data Fig. 5a). Strikingly, we observed a drastic decrease in phagocytosis of synaptosomes by primary mouse microglia cultured in high (5 mM) fructose compared to microglia cultured in control media or low (1 mM) fructose (**Fig. 2c,d**). Furthermore, we observed a significant decrease in phagocytosis of apoptotic neurons by primary mouse microglia cultured in high fructose compared to microglia cultured in control media (Extended Data Fig. 5c). These findings are important because even modest reductions in synaptic pruning or apoptotic neuron clearance can have catastrophic effects on neurodevelopment^9,44,87^.

**Fig. 2:**
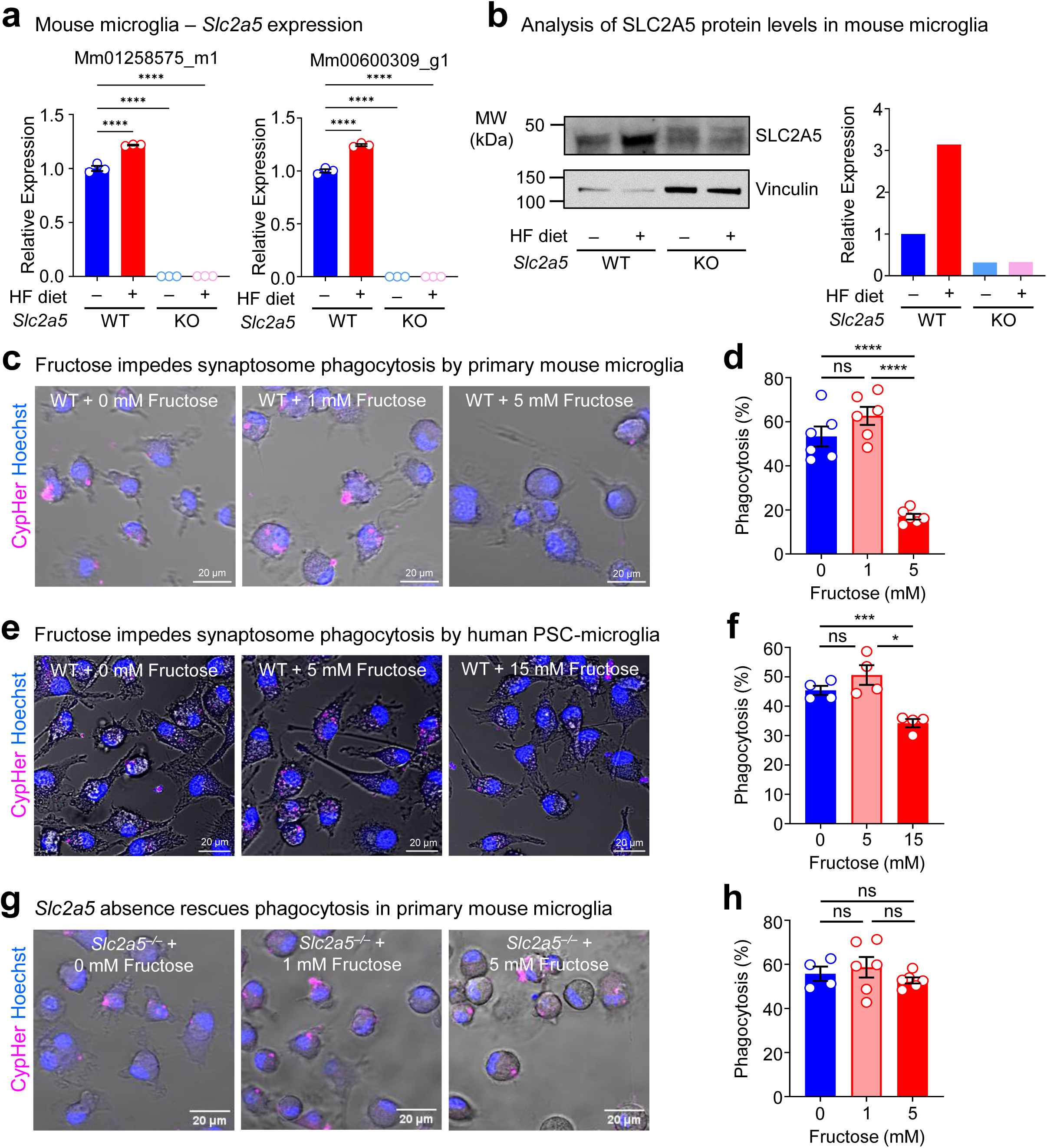
High fructose exposure directly suppresses microglia phagocytosis. **(a)** <u>High fructose exposure enhances *Slc2a5* expression in neonatal microglia.</u> Microglia from wildtype (WT) and *Slc2a5*^−/−^ (KO) P7 neonates born to and nursed by dams on control diet (CD) or high fructose diet (HF) as in **Fig. 1b** were isolated for analysis of *Slc2a5* expression via RT-qPCR. Shown is data from three different mice per condition using two different probes. Expression is normalized to 18s. **(b)** <u>High fructose exposure enhances SLC2A5 protein levels in neonatal microglia.</u> Experiments were performed as in **(a)** but instead processed for analysis via western blot. Shown is a representative image (left) and quantification (right) of SLC2A5 relative to vinculin. **(c, d)** <u>High fructose impedes phagocytosis of synaptosomes by primary mouse microglia.</u> Representative confocal microscopy images **(c)** and quantitation **(d)** of microglia phagocytosis of CypHer5E-labeled synaptosomes. CypHer5E+ microglia were identified in six different fields of view (FOVs) per condition and the percentage for each FOV is shown. See Extended Data Fig. 5a for detailed conditions and hexose concentrations. Data are presented as mean ± SEM and are representative of three independent experiments. **(e, f)** <u>High fructose impedes phagocytosis of synaptosomes by human pluripotent stem cell</u> <u>(PSC)-derived microglia.</u> Representative confocal microscopy images **(e)** and quantitation **(f)** of hPSC-derived microglia phagocytosis of CypHer5E-labeled synaptosomes. Cypher5E+ microglia were identified in four different FOVs per condition and the percentage for each FOV is shown. Data are presented as mean ± SEM and are representative of three independent experiments. **(g, h)** <u>GLUT5 absence rescues impaired phagocytosis of synaptosomes by primary mouse</u> <u>microglia.</u> Representative confocal microscopy images **(g)** and quantitation **(h)** of GLUT5-deficient microglia phagocytosis of CypHer5E-labeled synaptosomes. Cypher5E+ microglia were identified in at least four different FOVs per condition and the percentage for each FOV is shown. Data are presented as mean ± SEM and are representative of three independent experiments. Data are shown as shown as mean ± SEM. Significance was determined using one-way **(d, f, h)** or two-way ANOVA **(a)**. ns = not significant, \**p* < .05, \*\**p* < .01, \*\*\**p* < .001, \*\*\*\**p* < .0001. Statistical analyses were performed using the averages for each biological replicate, not pseudo-replications.

Next, we queried whether high fructose has a similar direct effect on human microglia. To test this, we derived microglia-like cells from human pluripotent stem cells (hPSCs)^88^ and cultured resultant cells in high fructose (15 mM), low fructose (5 mM), or control media. Similar to mouse primary macrophages, culturing hPSC-derived microglia in high fructose, but not low fructose, induced a significant increase in *SLC2A5* expression (Extended Data Fig. 5d). Importantly, hPSC-derived microglia cultured in high fructose exhibited significantly decreased phagocytosis of synaptosomes (**Fig. 2e,f**) and apoptotic neurons (Extended Data Fig. 5e), similar to mouse primary microglia cultured in high fructose. Taken together, high fructose directly suppresses phagocytosis capacity by both mouse and human microglia.

Finally, it is possible that the effect of high fructose on microglia is unrelated to fructose transport itself. To test this, we isolated primary microglia from GLUT5-deficient neonates and analyzed phagocytosis capacity under high fructose conditions. Importantly, GLUT5 deficiency completely reversed the decreased phagocytosis of synaptosomes (**Fig. 2g,h**) and apoptotic neurons (Extended Data Fig. 5f) observed in primary microglia cultured in high fructose. Thus, the phagocytosis-suppressive effect of high fructose on microglia is mediated by cell-intrinsic activity of the primary fructose transporter GLUT5.

### High fructose directly alters microglial metabolism

Whether fructose metabolism occurs in the CNS, especially in adults, remains an open question and may depend on a series of factors, including the technology used to measure fructose metabolism, what approach is used/which metabolites are measured, and the timing at which one looks. Given our finding that high fructose directly affects microglia phagocytosis, we sought to first determine if we could detect and quantify fructose metabolism in the neonatal brain. We turned to [2-^13^C]-fructose carbon nuclear magnetic resonance (NMR) because of its ability to rapidly and directly quantify fructose metabolism *in vivo*^89,90^. We injected [2-^13^C]-fructose into the intraperitoneal cavity of wildtype or GLUT5-deficient P7 neonates born to and nursed by dams on HF diet or control diet 15 min prior to analysis (**Fig. 3a**), which allowed us to accurately measure fructose metabolism in the CNS while limiting the potential confound of fructose conversion in the small intestine or liver^21^. Strikingly, we found increased catabolism of [2-^13^C]-fructose via fructolysis in the brains of wildtype neonates born to and nursed by dams on HF diet compared to wildtype neonates born to and nursed by dams on control diet ([2-^13^C]-lactic acid; **Fig. 3b**). This increased fructolysis was commensurate with an increase in the total lactate pool observed only in wildtype neonates born to and nursed by dams on HF diet (**Fig. 3c**). Importantly, both the increased catabolism of [2-^13^C]-fructose and the increased total lactate pool observed in wildtype neonates born to and nursed by dams on HF diet was completely reversed in neonates born to and nursed by dams on HF diet lacking GLUT5 (**Fig. 3b,c**). Thus, fructose catabolism (fructolysis) is observed in the brains of neonates on HF diet in a GLUT5-dependent manner.

**Fig. 3:**
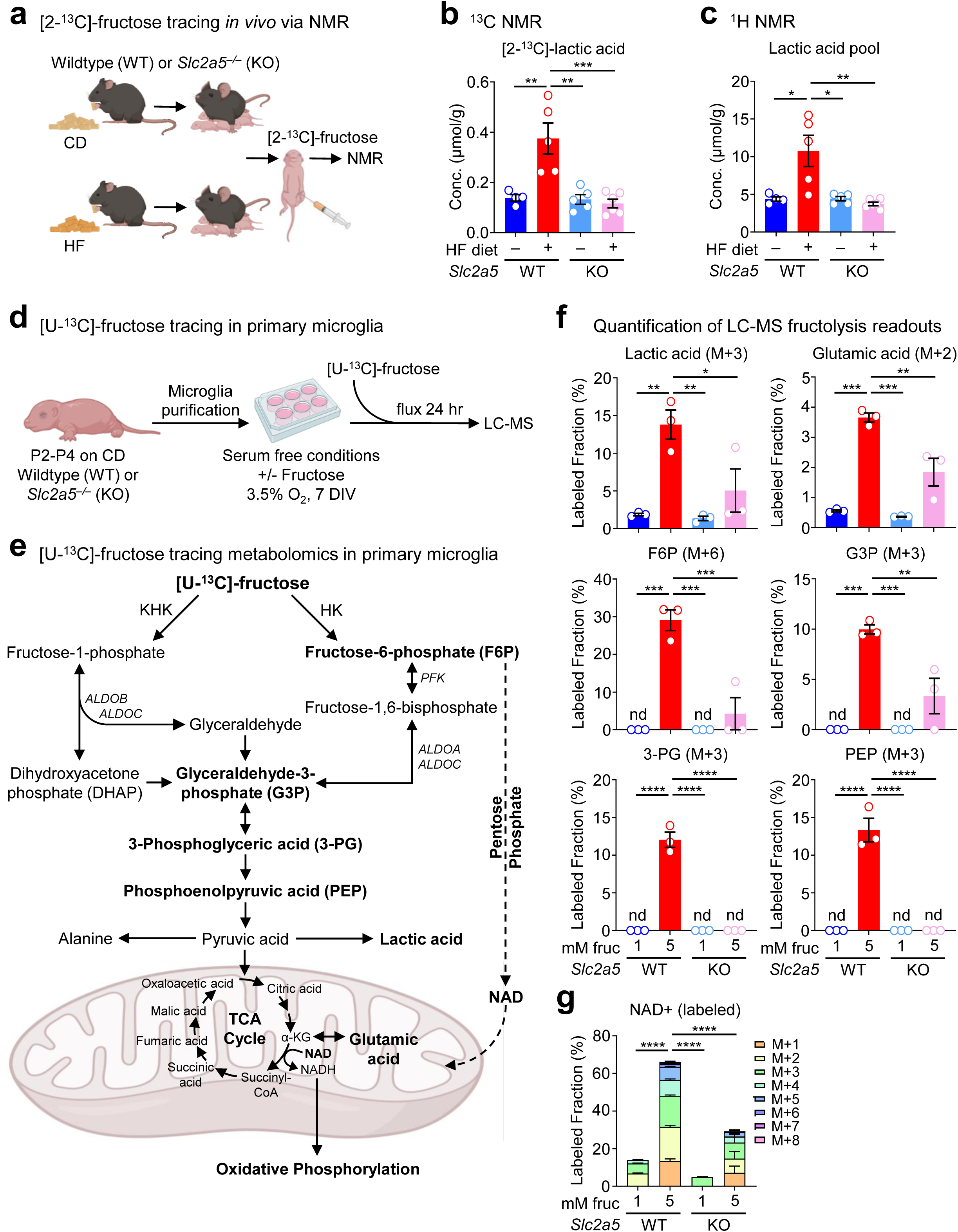
High fructose directly alters microglial metabolism. **(a-c)** <u>Neonatal brains exposed to HF exhibit increased fructolysis.</u> **(a)** Wildtype (WT) and *Slc2a5*^−/−^ (KO) P7 mice born to and nursed by dams on control diet (CD) or high fructose (HF) diet were injected intraperitoneally with 4 g/kg [2-^13^C]-fructose. Brains were isolated 15 min after infusion, immediately frozen, processed, and analyzed using proton (^1^H) and carbon (^13^C) NMR. **(b, c)** Quantitation of ^13^C-labeled (b) and total pool **(c)** of lactic acid in brains from P7 mice born to dams as outlined in **(a)**. Data are from four WT mice on CD (blue), five WT mice on HF (red), five KO mice on CD (light blue), and six KO mice on HF (pink) shown as mean ± SEM. Significance was determined by two-way ANOVA. \**p* < .05, \*\**p* < .01, \*\*\**p* < .001. **(d-g)** <u>Microglia treated with high fructose uniquely metabolize fructose via fructolysis.</u> **(d)** Schematic of experiment to assess fructose catabolism (fructolysis) in primary mouse microglia. Microglia were isolated from P2-P4 wildtype (WT) and *Slc2a5*^−/−^ (KO) mice nursed by dams on standard diet (SD) then cultured at physiological oxygen (3.5%) in serum-free conditions containing 5 mM glucose supplemented with either 1 mM or 5 mM fructose for one week (see Extended Data Fig. 5a). Each biological replicate is from microglia pooled from 4-5 littermates, with a total of three biological replicates per condition. On day 7, conditioning media was replaced with [U-^13^C]-fructose for 24 h at physiological oxygen, then cold-methanol extracted and analyzed using LC-MS. **(e)** Shown is the pathway of fructose catabolism (fructolysis). Intermediates in bold are metabolites detected by LC-MS. Hexokinase (HK, predominantly HK2 in microglia), phosphofructokinase (PFK), alpha-ketoglutarate (α-KG), aldolase (ALDOA, ALDOB, ALDOC), ketohexokinase (KHK). **(f, g)** Fractional enrichment of fructolysis metabolites including lactic acid, glutamic acid, fructose 6-phosphate (F6P), glyceraldehyde 3-phosphate (G3P), 3-phosphoglyceric acid (3-PG), phosphoenolpyruvic acid (PEP) **(f)**, and nicotinamide adenine dinucleotide (NAD+) **(g)**. See also Extended Data Fig. 6 for relative abundance of the identified metabolites. Data are shown as mean ± SEM. Significant was calculated by two-way ANOVA. \**p* < .05, \*\**p* < .01, \*\*\**p* < .001, \*\*\*\**p* < .0001.

We next sought to determine if microglia specifically catabolize fructose and how high fructose exposure affects microglia metabolism. To this end, we performed tracing of uniformly-labeled ^13^C [U-^13^C]-fructose in wildtype and GLUT5-deficient primary microglia conditioned in either low (1 mM) or high (5 mM) fructose in the presence of physiological glucose and oxygen (**Fig. 3d,e**). Wildtype microglia derived from neonates exposed to standard diet conditioned in both low and high fructose were capable of consuming and catabolizing fructose, demonstrated by ^13^C-labeled lactic acid (M+3) and glutamic acid (M+2; **Fig. 3f** and Extended Data Fig. 6a). However, consistent with our observation that microglia upregulate *Slc2a5* in response to high fructose, we observed that HF-conditioned microglia consumed more fructose (M+6 fructose; Extended Data Fig. 6a) and catabolized significantly more fructose as indicated by increased fractional enrichment and relative abundance of ^13^C-labeled lactic acid and glutamic acid (**Fig. 3f** and Extended Data Fig. 6a). Surprisingly, we found that HF-conditioned microglia exhibited drastic increases in both relative abundance and fractional enrichment of fructolysis intermediates (**Fig. 3f** and Extended Data Fig. 6b). This difference was particularly striking for the hexokinase (HK) product fructose 6-phosphate (F6P; **Fig. 3f** and Extended Data Fig. 6b). This is especially important because two recent reports demonstrate that deletion of the HK isoform HK2 results in increased microglia phagocytosis, whereas enforcing HK2 activity results in defective microglia phagocytosis^91,92^.

Although our data to this point suggest that fructose transport via GLUT5 is necessary for the detrimental effects of HF on microglia phagocytosis, it remains possible that the rescue observed in GLUT5-deficient microglia is independent of the metabolic changes observed in HF-treated microglia. However, consistent with our hypothesis that HF-induced alterations of microglia metabolism result in decreased phagocytosis capacity, we found that fructose uptake and catabolism in HF-treated, GLUT5-deficient microglia were reverted to levels near that of microglia conditioned in low fructose (**Fig. 3f** and Extended Data Fig. 6a). Additionally, the striking shift towards HK-mediated fructolysis observed in HF-conditioned microglia was almost completely reversed in HF-treated, GLUT5-deficient microglia (**Fig. 3f** and Extended Data Fig. 6b).

We and others recently reported that adult tissue-resident macrophages residing in prolonged physiological low oxygen are metabolically reprogrammed into a reduced state, using metabolic substrates to maintain redox homeostasis while performing efficient homeostatic apoptotic cell clearance^81,93^. Consistent with these findings, microglia conditioned in low fructose at physiological oxygen (3.5%) efficiently synthesized and accumulated reduced glutathione from fructose, whereas HF-conditioned microglia displayed a modest loss of ^13^C-labeled reduced glutathione while instead exhibiting significantly increased oxidized glutathione (Extended Data Fig. 6c). This change from a reduced state to an oxidized state has also been reported in aged microglia, termed lipid droplet-accumulating microglia (LDAM), which exhibit significantly decreased phagocytic capacity^94^. LDAMs also had a significantly increased NAD+/NADH ratio and transcriptional changes suggesting altered metabolic state^94^. Interestingly, we found that HF-conditioned microglia had increased fractional enrichment and relative abundance of fructose-derived NAD+ (**Fig. 3g** and Extended Data Fig. 6c) and altered contribution of fructose carbons into the TCA cycle (Extended Data Fig. 7). Importantly, however, the oxidative state observed in HF-treated microglia was significantly reversed in GLUT5-deficient microglia (**Fig. 3g** and Extended Data Fig. 6c, 7). Collectively, our data suggest that high fructose exposure, via GLUT5-dependent transport and fructose catabolism (**Fig. 3e**), alters microglia metabolism towards a potentially non-phagocytic metabolic state.

### Early life high fructose exposure contributes to cognitive defects and the development of anxiety-like behavior

Defects in microglial pruning, especially during neonatal development, can result in impaired learning and behavior^44,95^. Given the importance of microglial phagocytosis in the first week of life, we sought to determine whether excess early life fructose exposure directly causes cognitive defects, and whether those impairments were dependent on GLUT5. To address this, wildtype or GLUT5-deficient mice were raised by dams on CD or HF diet until weaning, then subjected to three key behavioral assessments previously shown to be affected by perturbed microglial phagocytosis: novel object recognition, modified Barnes maze, and fear extinction^96–99^. First, we performed novel object recognition (NOR; **Fig. 4a**), which is an experiment that broadly assesses animal cognition and recognition memory by applying the principle that mice prefer novel objects over objects previously introduced^100^. We found that mice born to and nursed by dams on HF exhibited no preference for the novel object (∼50%) compared to mice born to and nursed by dams on CD (∼75%, **Fig. 4b**). On the other hand, the loss of novel object preference observed in mice born to and nursed by dams on HF was completely reversed in GLUT5-deficient mice born to and nursed by dams on HF (**Fig. 4b**). Second, we performed the modified Barnes maze task (**Fig. 4c**), which specifically assesses spatial and working memory^101^. In contrast to our results with the NOR task, mice born to and nursed by dams on HF did not exhibit a significant defect in any parameters assessed, including the time to locate the escape hole (primary latency), time to enter the escape hole (total latency; **Fig. 4d**), number of holes incorrectly checked before locating the correct one (primary errors), and number of holes incorrectly checked before entering the correct one (Extended Data Fig. 8). Instead, mice born to and nursed by dams on HF displayed modestly better performance across some metrics that was reversed in GLUT5-deficient mice (**Fig. 4d** and Extended Data Fig. 8). The contradiction between NOR and Barnes maze performance suggests that HF affects cognition (assessed by NOR) and not memory (assessed by both NOR and Barnes maze). Finally, we performed fear extinction (**Fig. 4e**), which is an associative learning task that tests both the acquisition of fear through Pavlovian learning (conditioning) and the subsequent reversal of that acquired fear (extinction)^102^. This test is particularly important because deficits in fear extinction after an environmental threat has passed have been implicated in various anxiety disorders including PTSD^102^. Consistent with our observation that mice born to and nursed by dams on HF do not have altered memory, an equivalent number of mice acquired a fear response across conditions. Strikingly, however, mice born to and nursed by dams on HF exhibited drastically impaired fear extinction compared to mice exposed to CD (**Fig. 4f**). Importantly, the deficits in fear extinction observed in mice born to and nursed by dams on HF were completely reversed in GLUT5-deficient mice born to and nursed by dams on HF (**Fig. 4f**). Thus, exposure to fructose in early life contributes to cognitive defects and the development of anxiety-like behavior that is rescued by GLUT5 deficiency.

**Fig. 4:**
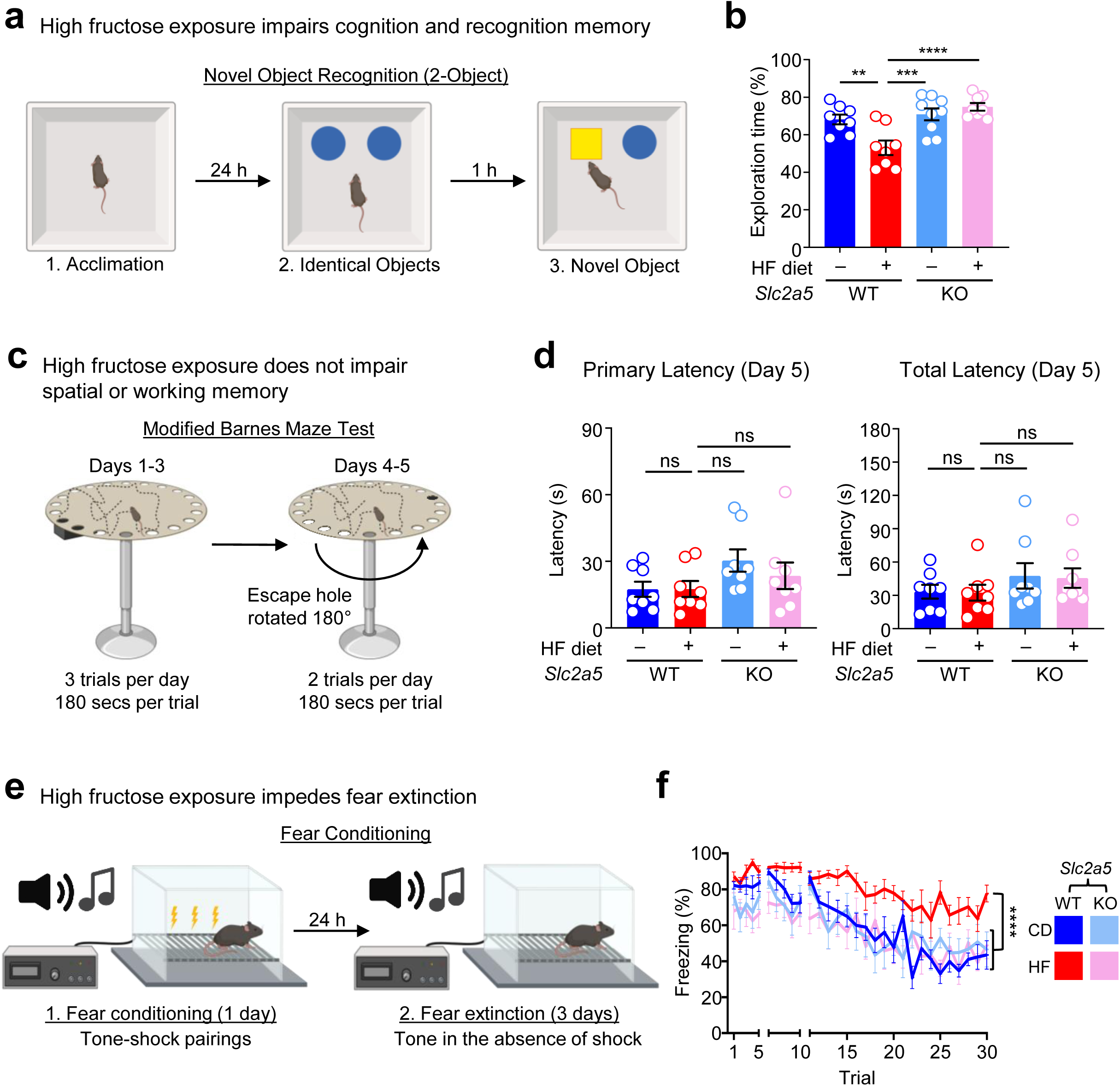
Early-life high fructose exposure impairs juvenile behavioral adaptability. **(a, b)** <u>High fructose exposure impairs cognition and recognition memory.</u> (a) Schematic of the novel object recognition (NOR, 2-object) task used to measure animal cognition and recognition memory. Mice were first acclimated to an empty arena and allowed to explore for 5 min. The following day, mice were placed in the same arena in the presence of two identical objects and allowed to explore for 5 min. After an inter-session interval of 1 h, one of the two original objects was replaced with a novel object and mice were allowed to explore for another 5 min. A blinded investigator quantified the amount of time spent exploring the novel object as a percentage of the total exploration time. **(b)** Offspring from wildtype (WT) and *Slc2a5*^−/−^ (KO) mice were raised by dams on control diet (CD) or high fructose diet (HF) until weaning. Weaned juveniles (age range: P26-P34) were subsequently tested as described in **(a)**. Data are from eight WT on CD (blue), eight WT on HF (red), nine KO on CD (light blue), and eight KO on HF (pink), shown as mean ± SEM. Significance was determined using two-way ANOVA. \*\**p* < .01, \*\*\**p* < .001, \*\*\*\**p* < .0001. **(c, d)** <u>Evaluation of spatial learning and memory via the modified Barnes Maze</u>. Experiments were performed as in **(b)**, but mice were used to assess spatial learning and memory. **(c)** Schematic of experiments used to analyze spatial learning and memory using the modified Barnes Maze. After three training days during which the escape hole was placed at the same location for three trials each day, the escape hole was then moved 180° on days 4 and 5 and two trials per day were performed. **(d)** Each of the four cohorts in **(b)** were tested as outlined in **(c)**. Shown is the time to locate (left, primary latency expressed in seconds) and enter (right, total latency expressed in seconds) the escape hole. Data are shown as mean ± SEM. Significance was determined using two-way ANOVA. ns = not significant. **(e, f)** <u>High fructose exposure impedes fear extinction.</u> Experiments were performed as in **(b)**, but mice were lastly used to assess fear extinction. **(e)** Schematic of experiments used to analyze fear extinction. Mice were exposed to five trials of paired tone and shock to condition fear response (freezing) in response to tone. After 24 h, mice were moved to a new contextual environment (extinction chamber) and exposed to five consecutive tone presentations without paired shock on day 1 (trial 1-5) and day 2 (trial 6-10) then twenty trials on day 3 (11-30) to assess maximum extinction. **(f)** Each of the four cohorts in **(b)** were tested as outlined in **(e)**. Shown is the percentage of time mice freeze in response to tone. Data are shown as mean ± SEM. Significance was determined using two-way ANOVA of the area under the curve (AUC) for each cohort. \*\*\*\**p* < .0001.

## Discussion

The chronic excessive consumption of fructose has contributed to a major public health crisis because of its contribution to the increased prevalence of metabolic disease. Recent epidemiological evidence suggests that the negative effects of high fructose diet may extend beyond metabolic disease and be associated with the onset of mood and anxiety disorders especially in adolescents. Here, we find that high fructose diet, delivered either during pregnancy or directly to early perinatal mice, dramatically decreases microglia phagocytic potential including disrupted synaptic pruning and clearance of apoptotic neurons. We show that fructose acts directly on both mouse and human microglia by driving microglia metabolism towards a metabolic state that suppresses phagocytosis. Importantly, deletion of SLC2A5 (GLUT5), the primary fructose transporter uniquely expressed in microglia, rescued the altered metabolism and phagocytosis deficiency caused by high fructose. Finally, we show that mice born to dams on high fructose diet exhibit cognitive deficits and develop anxiety-like behaviors that are reversed in neonates lacking GLUT5. Taken together, our work provides a mechanistic explanation for the epidemiological observation that exposure to high fructose early in life is associated with increased anxiety disorder prevalence in adolescents.

Our work revealed that exposing microglia to high fructose drives the catabolism of fructose into fructose 6-phosphate. Fructose is known to be catabolized via two different enzymes: ketohexokinase into fructose 1-phosphate, which is dominant in the liver and the intestine, or hexokinase into fructose 6-phosphate. Interestingly, two recent studies found that hexokinase 2 (HK2), the main isoform in microglia, is a negative regulator of microglia phagocytosis^91,92^. Specifically, both groups demonstrated that deletion or pharmacological inhibition of HK2 caused increased microglia phagocytosis activity. On the other hand, supplementation with glucose 6-phosphate or fructose 6-phosphate, but not fructose 1,6-bisphosphate, suppressed microglia phagocytosis activity^91^. These findings suggest that HK2 enzymatic activity directly impedes microglial phagocytosis, consistent with our finding that microglia exposed to HF display increased synthesis of fructose 6-phosphate from fructose and dramatically decreased phagocytic activity. Our findings, and the work on HK2, are particularly interesting when considering that mitochondrial metabolic activity is enhanced in early neonatal mice (P9) and is required for microglia phagocytosis, but subsides after the critical microglia phagocytic window^103^. Future work will be required to determine how hexose metabolism, HK2 enzymatic activity, and mitochondrial metabolism specifically function to support or suppress neonatal microglia phagocytosis.

Although microglia uniquely express SLC2A5 (GLUT5), its expression increases in both mouse and human microglia during aging. It remains unclear what, if any, homeostatic function GLUT5/fructose transport and metabolism serve in adult microglia. Here, we show that neonatal microglia use GLUT5 to import and metabolize fructose, but doing so suppresses capacity to perform phagocytic functions essential during development. Given that microglia transition from a highly phagocytic state during the critical neonatal window to a quiescent, non-phagocytic state in adulthood, we speculate that GLUT5 upregulation, and concomitant increased fructose import/catabolism, is determinative of this transition and supports microglial longevity. This hypothesis is supported by multiple observations. For instance, microglia are metabolically flexible, capable of surviving in an environment that preferentially delivers glucose to astrocytes and neurons^104^. Additionally, microglia reside in physiologically low oxygen often for the entire life of the host^83–85^. We and others recently reported that peripheral tissue-resident macrophages residing in physiological hypoxia downregulate total glucose consumption and catabolism, instead efficiently shunting glucose into a noncanonical pentose phosphate pathway loop that supports redox homeostasis^81,93^. Whether increased GLUT5 expression allows microglia to better use the ∼200 μM fructose present in the human adult CNS^105^ and if this goes awry in unhealthy aging (e.g., neurodegeneration) remains unknown. Understanding how adult microglia use GLUT5 and fructose in health and disease will be an important area of work moving forward, particularly given the prevalent use of high fructose corn syrup as an additive in the common Western diet.

Our findings that high fructose exposure results in defects in the novel object recognition (NOR) task and fear extinction but not the modified Barnes maze suggest that early life fructose is specifically affecting animal cognition and behavior (e.g., the development of anxiety-like behaviors) but not memory. This is consistent with extensive prior work suggesting that fear extinction is not simply a matter of ‘forgetting’ the previously learned fear^106^. However, a previous report suggests that, in adult mice, microglia are also important for ‘forgetting’ of previously learned contextual fear memories via activity-dependent synaptic pruning^97^. Thus, it remains possible that high fructose exposure may also affect microglia function in the adult CNS, albeit by preventing microglia synapse elimination and removal of memory engrams.

Finally, our results have potentially profound implications not only for pregnancy (via transfer of fructose via breast milk^4,49^) but also adolescent development. In humans, microglial synaptic pruning and phagocytosis continues well into the second decade of life^107,108^ potentially broadening the window when high fructose consumption could be detrimental. High fructose consumption itself likely does not cause the development of mood or anxiety disorders. However, it is plausible that high fructose consumption serves as the first ‘hit’, while a subsequent trauma is required to trigger the development of a disorder^109–111^. Nevertheless, we show for the first time that a common constituent of Western diet, high fructose, directly impacts microglia phagocytosis and neonatal neurodevelopment. How this factors into human adolescent development itself, given the recent rise in mood and anxiety disorder development, especially post-COVID19 pandemic, remains an important public health question.

## Materials and Methods

### Contact for Reagent and Resource Sharing

For further information and requests for resources and reagents should be directed to the Lead Contact, Justin S. A. Perry.

### Mice

Wild-type C57BL/6J mice (Stock No. 000664) were purchased from The Jackson Laboratory. *Slc2a5*^−/−^ (KO) mice were originally generated by Wu et al.^79^ and provided by M. Zeng. For experiments using standard diet (SD), dams were provided PicoLab Rodent Diet 5053 (LabDiet 5053, PMI). All mice were housed at Memorial Sloan Kettering Cancer Center (MSKCC) or Weill Cornell Medicine under specific pathogen-free (SPF) conditions with 12-hour light/dark cycles and were cared for by Research Animal Resource Center (RARC). Mice were housed under ambient conditions and provided *ad libitum* access to water and food. All studies conducted were approved by the Sloan Kettering Institute (SKI) and Weill Cornell Medicine Institutional Animal Care and Use Committee (IACUC).

### Early life high fructose diet and delivery

Isocaloric control and high fructose diets are modified forms of the AIN-93G Formula generated by Research Diets, Inc. Control diet (CD, D19060506) contains 0 kcal% sucrose and high fructose diet (HF, D19060507) contains 15 kcal% fructose. Dams of each cohort (CD and HF) were placed on respective diets at least one week before mating and diets were maintained throughout gestation and lactation. For intragastric delivery of fructose, mice received 50 μL of sterile water with or without fructose at 45 mg/mouse. Solutions were delivered daily via injections into the visible milk spot of wildtype (WT) or *Slc2a5*^−/−^ neonates from perinatal day (P)1 to P7. The selected fructose dosage is 45% of the typical 100 mg/mouse dose delivered daily by oral gavage to adult mice which was determined to be equivalent to 3% of total daily caloric intake and analogous to human consumption of less than 12 oz of sugar sweetened beverage^112^.

### Immunofluorescence and confocal microscopy

Neonatal mice were anesthetized with CO_2_, perfused with 10 mL cold PBS (Corning, 21-040-CM), and harvested brains were submerged in 4% formaldehyde (StatLab, 28530-1) for 24-48 hours at 4°C. Tissues were then washed 3x in PBS, embedded in 2% SeaPlaque Agarose (Lonza, 50101), and sectioned by vibratome (Leica, VT1000S) into 50 μm sections. Sections were placed in 12 or 24 well plates for permeabilization using 1% Triton X-100 (Sigma-Aldrich, T9284) and 5% BSA (Sigma-Aldrich, A7888) in PBS overnight at 4°C with rotation. After 3x PBS washes, free floating sections were blocked with 5% normal goat serum (Jackson ImmunoResearch, 005-000-121) and 0.1% Triton X-100 in PBS (blocking buffer) for 1 hour at room temperature. Sections were incubated for 48-72 hours with the following primary antibodies at 1:400: rabbit anti-Iba1 (Wako, 019-19741) and mouse anti-PSD-95 (Sigma-Aldrich, MAB1598). Sections were then washed 3x with PBS containing 0.05% Tween 20 and incubated with Alexa Fluorophore conjugated goat anti-rabbit IgG (H+L) secondary antibody Alexa Fluor 488 (Thermo Fisher, A-11034) and goat anti-mouse Alexa Fluor Plus 647 (Thermo Fisher, A32728) at 1:400 in blocking buffer covered at room temperature for 2 hours with rotation. Sections were washed again 3x with PBS containing 0.05% Tween 20 (Sigma-Aldrich, P1379) but with the addition of 1:2000 Hoechst 33342 (Thermo Fisher, H3570) during the second wash before mounting with ProLong Gold antifade reagent (Thermo Fisher, P36934). For TUNEL staining, manufacturer’s instructions for the In Situ Cell Death Detection Kit, TMR red (Roche, 12156792910) were followed before mounting. Images of neonatal brain sections were taken as 20 μm Z-stacks on a Zeiss LSM 980 with Airyscan 2 (20X objective). For each mouse, a minimum of 4 fields of view (FOV) were obtained from the prefrontal cortex and cortex before maximum intensity z projections were created in Fiji for manual quantification and automated morphological analysis. Max intensity z projections were compiled from 20X images to best capture larger population level effects.

### Analysis of microglia morphology

Iba1-labeled microglia were converted to gray scale using the MorphoLibJ plugin in Fiji (opening filter, area minimum: 25 pixels, connectivity: 8) and aligned to an octagon morphological filter element (radius pixels: 1) as previously described^113^. Threshold detection levels were adjusted to align automated microglia counts with previously obtained manual counts to ensure optimal detection of complete cell somas. Microglia were analyzed using the Analyze Particles function with a size limit of 10 pixels to exclude background noise. Major axis length was derived from the longest axis of an ellipse fit to each microglia soma. A combination of major axis length and soma area were used to compute a cell roundness score per cell (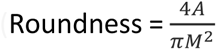, A = soma area, M = major axis length)^113^, and then cell roundness values were averaged per field of view (4 FOVs per animal). For analysis of ramification and branching, identical images were re-imported into Fiji and filtered by the Unsharp Mask function (radius: 3 pixels). A new threshold was obtained to capture branching and junctions. The Close binary function was applied and the binary skeletonized, all as previously outlined^114^. Once skeletonized, despeckling and outlier removal were applied as needed and branching metrics exported for analysis (Extended Data Fig. 1a-b).

### Mixed glial cultures

Meninges were removed from cortices harvested from WT P2-P4 neonates in cold HBSS (Thermo Fisher, 14025092). The dissected cortices were then manually dissociated in HBSS and spun at 350 G for 5 min. The homogenate was resuspended in 10 mL warm DMEM (Corning, 10-017-CV) supplemented with 10% (vol/vol) heat-inactivated FBS (Sigma, 12306C), 100 U/mL of penicillin and 100 μg/mL streptomycin (Gibco, 15140-122), and 2 mM L-glutamine. Cortices from two neonates were cultured in each Poly-D-Lysine (Thermo Fisher A3890401) coated T25 flask with 10 mL DMEM with or without 5 mM fructose, with osmolarity controlled by mannitol (Sigma-Aldrich, M4125). Mixed glial cultures were grown in a standard incubator in humidified 5% CO_2_ at 37°C. Media was changed 2 days later and then every 3-4 days. After 18 days, microglia were shaken off at 200 rpm at 37 °C for 1 hour and quantified per flask using the Countess II (Thermo Fisher) to obtain mixed glial culture yield for both control and fructose treated flasks.

### Primary microglia isolation and culture

Neonatal brains were harvested from P2-P4 WT and *Slc2a5*^−/−^ mice and dissociated using the Miltenyi gentleMACS Octo Dissociator with Heaters, brain dissociation kits (Miltenyi Biotec 130-096-427, 130-107-677, 130-092-628), and gentleMACS C Tubes (Miltenyi Biotech 130-093-237), following manufacturer’s instructions. Microglia were purified using CD11b MicroBeads (Miltenyi Biotec 130-097-142), LS Columns (Miltenyi Biotech 130-042-401) and the QuadroMACS separator (Miltenyi Biotech 130-091-051), following manufacturer’s instructions. Isolated primary microglia were then cultured in previously established serum-free TIC media^86^ containing key factors including TGF-β2, IL34, and cholesterol, but modified to use DMEM without glucose (Gibco, A14430-01) to create specific conditions with various glucose (Sigma-Aldrich, 68270) and fructose (Sigma-Aldrich, F0127) concentrations along with the nonmetabolizable sugar mannitol to control for osmolarity. For *in vitro* conditioning, microglia were grown in standard tissue culture incubator conditions, except with 3.5% O_2_ in the BioSpherix X3 Xvivo System.

### Primary microglia phagocytosis assays

For all experiments, microglia from P2-P4 WT and *Slc2a5*^−/−^ mice were isolated and incubated as described above with serum free media conditions containing indicated hexose combinations. For synaptosome phagocytosis assays, synaptosomes were first purified from WT adult mice using the Syn-PER Synaptic Protein Extraction Reagent according to manufacturer’s instructions (Thermo Scientific, 87793). Synaptosomes were then stained with CypHer5E (GE Life Sciences, PA15401) via agitation, washing, and sonication. A BCA assay was used to determine the staining CypHer5E concentration (10 mM CypHer5E/50 uL). CypHer5E+ synaptosomes were added to each chambered well (Nunc, 155409) conditioned at 3.5% O_2_ with hexose concentrations corresponding to the growth condition for each well (0 mM, 1 mM, or 5 mM fructose with constant 5 mM glucose, and 20 mM, 19 mM, or 15 mM mannitol, respectively, to control for osmolarity). Phagocytosis was assessed using time-lapse confocal microscopy on a Zeiss Axio Observer Z1-7 inverted fluorescence microscope fully encased in a black-out environmental chamber, equipped with a Z PIEZO stage encapsulated with a heating/gas-controlled insert, a Zeiss 20x PlanApo (0.8 NA) objective, a 6-channel, 7-laser LSM 980 (405nm, 445nm, 488nm, 514nm, 561nm, 594nm, 639nm), and a Airyscan 2 multiplex detector. Chambered glass slides containing cells were removed from 3.5% O_2_ at appropriate time intervals so that images for each condition could be immediately taken at the same time point. For each condition, four regions containing similar cell numbers were selected and imaged. Data was acquired using Zen Blue software (Zeiss). For neuron efferocytosis assays, apoptosis was induced in the Neuro-2a (N2A) neuroblastoma cell line (ATCC, CCL-131) using 0.25 μM staurosporine (Cayman Chemical, 81590) for 12-14 hours. Primary microglia were isolated, spot plated (50,000 cells in 50 μL) in 24 well plates and conditioned for one week as described above. Microglia were incubated with target cells at a 1:1 phagocyte:target ratio for 30 min at 3.5% O_2_, scraped off wells, and CypHer5E+ microglia were analyzed using an Attune NxT flow cytometer (Thermo Fisher). Samples were analyzed with FlowJo v10.9 (BD). When primary microglia are used as phagocytes, there is an inherent difference in the absolute percentage uptake of corpses between experiments performed on different days. Therefore, phagocytic index was used to compile data from multiple experiments. Phagocytic index = percent engulfment (experimental/control).

### Human pluripotent stem cell-derived microglia generation

Human pluripotent stem cells (PSCs) were generated from anonymized human PBMCs based on published protocols^115^ using Sendai viral vectors (Thermo Fisher, A16517) driving high expression of pluripotency markers NANOG and OCT4, which were confirmed by flow cytometry. PSCs were maintained on CF1 mouse embryonic fibroblasts (Thermo Fisher, A34181) in Embryonic Stem Cell (ESC) medium (Thermo Fisher, KO-DMEM 10829-018; 20% KO serum replacement 10828-028; 2 mM L-glutamine 25030-024; 1% nonessential amino acids 11140-035; 1% penicillin/streptomycin 15140163; 0.2% β-mercaptoethanol 31350-010) supplemented with 10 ng/mL basic fibroblast growth factor bFGF (PeproTech, 100-18B). The strategy for PSC hematopoietic differentiation was adapted from a previously published protocol^116^ in which embryoid bodies (EBs) were formed by dissociating and maintaining PSCs in ESC media supplemented with 10 μM Rho-associated protein kinase (ROCK) inhibitor (Sigma, Y0503) while being kept on an orbital shaker at 100 rpm for 6 days to allow for spontaneous formation of EBs with hematopoietic potential. At day 6 of differentiation, 200-500 μm cystic EBs were selected under a dissecting microscope and transferred onto adherent tissue culture plates (∼2.5 EBs/cm^2^) for cultivation in Hematopoietic Differentiation (HD) medium (APEL 2, Stem Cell Tech, 05270; Protein Free Hybridoma, Thermo Fisher, 12040077; penicillin and streptomycin, Thermo Fisher, 15140163) from day 6 up to day 18 from the start of differentiation, supplemented with 25 ng/mL human IL-3 (Peprotech, 200-03) and 50 ng/mL human M-CSF (Peprotech, 300-25). At day 18 of differentiation, macrophages produced by EBs were collected from suspension and cultivated on tissue culture plates at a density of ∼15,000 cells/cm^2^ in RPMI (Thermo Fisher, 61870036) supplemented with 10% FBS and human M-CSF (100 ng/mL) for 6 days before use for downstream experiments. Engulfment assays using PSC derived cells were performed as described above for primary microglia, except cells were plated at 15,000/cm^2^ in 24 well plates with constant 10 mM glucose plus 0 mM fructose and 15 mM mannitol (to control for osmolarity), 5 mM fructose and 10 mM mannitol, or 15 mM fructose and 0 mM mannitol.

### [2-^13^C]-fructose NMR

WT and *Slc2a5*^−/−^ P7 neonatal mice born to dams on HF or CD were injected intraperitoneally via the quadriceps with 4 g/kg [2-^13^C]-fructose (Cambridge Isotope Laboratories, CLM-1527) in PBS (20 mg in 200 μL PBS). Brains were harvested and flash frozen in liquid nitrogen. Sample extraction was performed using methanol (Sigma-Aldrich, 34860), chloroform (Sigma-Aldrich, 366927), beads (Fisherbrand 15-340-151), and a Bead Mill (Fisherbrand 15-340-163). Brain extracts were spun in a centrifugal evaporator (GeneVac) for 5 hours, and pellets were dissolved in 600 μL of 1 mM TSP, 10 mM imidazole, and 0.2 wt% sodium azide in deuterium oxide. After sonication for 30 minutes, the solutions were vortexed and added to NMR tubes. Carbon and proton NMR was performed using 1 mM TSP as a chemical shift and concentration internal standard.

### [U-^13^C]-fructose tracing metabolomics

Primary microglia were isolated as described above using CD11b microbeads. One million cells were pooled from 4-5 P2-P4 neonates for each of the three biological replicates per condition. Cells were cultured for 1 week at 3.5% O_2_ in 1 mM or 5 mM fructose plus constant 5 mM glucose in TIC media, with a media change after 4 days. On the final day all media conditions were replaced with 5 mM [U-^13^C]-fructose (Cambridge Isotope Laboratories, CLM-1553). Metabolites were extracted using 80% methanol 24 hours later, on dry ice. The supernatants containing polar metabolites were dried down and dissolved in water. Targeted LC-MS analyses were performed on a Q Exactive Orbitrap mass spectrometer (Thermo Fisher) in polarity-switching mode, coupled to a Vanquish UPLC system (Thermo Fisher). A Sequant ZIC-HILIC column (2.1 mm i.d. × 150 mm, Merck) was used for separation of metabolites. Flow rate was set to 150 μL/min. Buffers consisted of 100% acetonitrile for mobile B, and 0.1% NH_4_OH/20 mM CH_3_COONH_4_ in water for mobile A. Gradient was from 85% to 30% B in 20 min followed by a wash with 30% B and re-equilibration at 85% B. Data analysis was performed using El-MAVEN (v0.12.0). Metabolites and their ^13^C isotopologues were identified based on exact mass within 5 ppm and standard retention times. Relative metabolite quantitation was performed based on peak area for each metabolite.

### Novel object recognition

The novel object recognition (NOR) task was conducted as previously described^117,118^. Briefly, all tests were conducted in a plastic box measuring 29 cm × 47 cm × 30 cm, with stimuli consisting of similarly sized plastic objects that varied in color and shape. A camera mounted directly above the box recorded all testing sessions for video analysis. One day prior to testing, mice were acclimated to the room and chamber, with 30 min in the testing room and 5 min to explore the empty arena. Mice were placed in the same arena 24 hours later in the presence of two identical objects and were allowed to explore for 5 min. After an intersession interval of 1 hour, mice were placed in the same box, with one of the two objects replaced by a novel object, and were allowed to explore for 5 min. Exploratory behavior was assessed manually, and exploration of an object was defined as a mouse sniffing the object or touching the object while looking at it. Placing the forepaws on the objects was considered as exploratory behavior, but climbing on the objects was not. Minimal exploration time (10 sec) for both objects during the test phase was used, and no significant difference in total exploration time was detected. The amount of time taken to explore the novel object was expressed as a percentage of the total exploration time and used to provide an index of recognition memory. Any-maze v7.1 (Stoelting) was used for acquisition and analysis of data.

### Barnes Maze

The Barnes maze test was performed as previously described^117,118^. Briefly, the maze consists of a circular open surface (90 cm in diameter) elevated to 90 cm and 20 circular holes (5 cm in diameter) equally spaced around the perimeter that were positioned 2.5 cm from the edge. A wooden escape box (11 × 6 × 5 cm) was positioned beneath one of the holes, and aversive stimuli included neon lamps and a buzzer. Extra-maze visual cues consisted of objects within the room including a table, computer, sink, door, plus the experimenter. No wall or intra-maze visual cues were placed around the edge of the maze. The ANY-maze tracking system (Stoelting) was used to record the movement of mice on the maze. Between trials, mice were placed into cages in a dark room adjacent to the test room for the intertrial interval (20-30 min). The acquisition phase consisted of three consecutive training days (days 1-3), with three trials per day during which the escape hole was located at the same location. For each trial, a mouse was placed into a start tube located in the center of the maze, the start tube was raised, and the buzzer was turned on until the mouse entered the escape hole. Between trials, the maze and escape box were cleaned with 10% ethanol in water to minimize olfactory cues. Mice were given 3 min to locate the escape hole for each trial and were guided to the escape box if they failed to enter the escape hole. Parameters of learning performance that were recorded included: latency to locate the escape hole (primary latency), latency to enter the escape hole (total latency), the number of errors made before locating the escape hole (primary errors), and the number of errors made before entering the escape hole (total errors). A mouse dipping its head into any hole not containing the escape box was considered an error. On days 4 and 5, the location of the escape hole was moved 180° from its previous location and two trials per day were performed.

### Fear conditioning and extinction

Fear conditioning and extinction assays were performed as previously described^119^. Briefly, mice were placed in shock chambers (Coulbourn Instruments) and were fear conditioned after 2 min of habituation with 3 tone-shock pairings consisting of a 30 sec (5 kHz, 70 dB) tone (conditioned stimulus, CS) that co-terminated with a 1 sec (0.7 mA) foot shock (unconditioned stimulus, US). The intertrial intervals (ITIs) between each tone-shock pairing were 30 sec. Mice remained in the conditioning chambers after the final tone-shock pairing for 1 min. For classical fear extinction with one session per day for 3 days, 24 hours after fear conditioning, mice were placed in extinction chambers, habituated for 2 min, and then exposed to 5 presentations of the tone (CS) in the absence of the shock (US), with each tone lasting for 30 sec with an ITI of 30 sec (5 trials on days 1 and 2, and 20 trials on day 3 to confirm maximal extinction). Mice remained in the extinction chambers for 1 min after the final tone presentation. Experiments were performed using Graphic State software (Coulbourn instruments) and all mice were video recorded for analysis. Freezing behavior was scored using previously validated MATLAB code and percent time spent freezing (freezing %) was calculated by dividing the amount of time spent freezing during the 30 sec tone presentations by tone duration.

### Reverse transcription quantitative PCR

Total RNA was extracted from microglia after CD11b microbead isolation from P7 neonates born to and nursed by WT and *Slc2a5*^−/−^ dams using the NucleoSpin RNA kit (Macherey-Nagel, 740955.250). The QuantiTect Reverse Transcription Kit (Qiagen, 205314) was used to synthesize cDNA according to the manufacturer’s instruction. Quantitative gene expression for mouse and human genes was performed using TaqMan probes and TaqMan Fast Advanced Master Mix (Applied Biosystems/Thermo Fisher, 4444554) with a QuantStudio 6 Pro (Applied Biosystems). For murine *Slc2a5*, 10 commercially available probes collectively spanning all 14 exons of the gene were tested and quantified relative to 18S. Details for TaqMan probes are listed in Extended Data Fig. 4b (mouse) and Extended Data Fig. 5e (human).

### Western blot

Primary microglia were isolated as described above, and cell pellets containing 1 million cells each were lysed in RIPA buffer (Thermo Fisher, 89900) with protease and phosphatase inhibitors (Thermo Fisher, 78440). Lysates were passed through a syringe with an 18G needle, spun at max speed for 20 min, and quantified using the BCA protein assay (Thermo Fisher, 23227) according to manufacturer’s instructions. Lysates were not boiled prior to loading, and 40 μg of each protein lysate were resolved on 4-12% Bis-Tris gels (Thermo Fisher, NP0322BOX) under denaturing conditions. Proteins were blotted onto a nitrocellulose membrane (Thermo Fisher, LC2009) and probed with anti-SLC2A5 (Santa Cruz, sc-271055) and anti-Vinculin (Sigma Aldrich, V9264) antibodies. Membranes were visualized using an anti-mouse HRP-conjugated secondary antibody (Cytiva, NA931), ECL reagent (RPN2209) and film (Fisher Scientific, NC9556985). Specific bands were quantified using Fiji with normalization to loading controls.

### Statistics and reproducibility

Data were analyzed using GraphPad Prism 10.0.0. Significance was determined using one of the following: unpaired two-tailed Student’s test (independent samples t-test), one-way ANOVA, or two-way ANOVA. Microglia from 4-5 P2-P4 neonates of the same genotype were pooled in independent experiments in order to generate sufficient material for U-^13^C fructose tracing and phagocytosis assays. Otherwise, single biologically-independent samples were included in statistical and graphical analyses. A minimum of 4 fields of view (FOVs) per animal or condition were analyzed by confocal microscopy. For analysis of microglia density and morphology, 4 FOVs per mouse were plotted as dots to represent the absolute number of cells counted or analyzed. For all other analysis, data from 4 FOVs per mouse were first averaged and then plotted as one dot representing one animal. In all cases, statistical analyses were performed using the averages for each animal, not pseudo-replications.

## Acknowledgements

We thank members of the Perry laboratory and co-authors for critical reading of this manuscript. We also thank the Weill Cornell Medicine Proteomics and Metabolomics Core Facility for LC-MS experiments and the Memorial Sloan Kettering Nuclear Magnetic Resonance Core for NMR experiments. This work was supported by grants to J.S.A.P. from the NIH (NCI 5R00CA237728; NIGMS 1DP2GM146337), a Pew Biomedical Scholars Award, a grant to K.R.K. from the NIH (NCI 1R01CA248364), a grant to M.Z. from the NIH (NICHD 1R01HD110118), training grant support to Z.W. (NIAID 5T32AI134632), A.T. (NCI 5T32CA009149), and T.S. (5T32CA254875-03), and MSKCC Cancer Center Support Grant P30CA008748. This work benefitted from data assembled by the ImmGen consortium. Some figure panels were created using the commercial version of BioRender.

## Contributions

Z.W. and J.S.A.P. planned, with Z.W. executing, the majority of experiments that A.L. assisted with. Z.L., A.J.T., I.C.M., T.S., W.S.R., P.S., J.E.R.P., and A.B. assisted with some *in vivo/in vitro* experiments. T.L. and F.G. assisted with human PSC experiments. C.M.T. and K.R.K. assisted with NMR experiments. G.F., L.G., C.D.L., C.N.P., and M.Y.Z. assisted with analysis and interpretation of *in vivo*/*in vitro* experiments. J.I.E. and C.D.L. provided important technical and theoretical discussions. K. R. K., and J.S.A.P. were responsible for the conceptualization of the project. All authors assisted in the preparation and review of the manuscript.

