## Supplemental Data for "Early life high fructose exposure disrupts microglia function and impedes neurodevelopment"

**Extended Data Fig. 1.**

**(a-d)** Intragastric delivery of fructose to neonates leads to decreased microglia numbers and increased uncleared apoptotic cells. **(a)** Neonatal wildtype (WT) and *Slc2a5*$^{-/-}$ (KO) mice were injected daily into the visible milk spot from P1 to P7 with fructose (45 mg per mouse) or sterile water control. Brains were then collected and analyzed by confocal microscopy. **(b)** Microglial cell count was quantified from four fields of view (FOVs) per mouse for each condition from **(a)** and number of microglia per FOV is plotted. Data are from six WT mice i.g. control (blue), six WT mice i.g. fructose (red), three KO mice i.g. control (light blue), and five KO mice i.g. fructose (pink). Data are shown as mean ± SEM. Significance was determined by two-way ANOVA. $^{*}p < .05$, $^{***}p < .001$. **(c)** Representative images and **(d)** quantitation of uncleared (free) apoptotic cells (pyknotic nuclei) from **(a)**, determined using Iba1 staining of microglia and Hoechst staining of nuclei. Arrows denote condensed chromatin (pyknotic nuclei) which is a hallmark indicator of apoptosis. Scale bars, 100 µm (inlay, 20 µm). **(d)** Quantification of average number of free pyknotic nuclei in **(c)**, from four FOVs per mouse with the average of those FOVs plotted per mouse. $n = 6$ (WT i.g. control), $n = 6$ (WT i.g. fructose), $n = 3$ (KO i.g. control), $n = 5$ (KO i.g. fructose). Data are from six WT mice i.g. control (blue), six WT mice i.g. fructose (red), three KO mice i.g. control (light blue), and five KO mice i.g. fructose (pink). Data are shown as mean ± SEM. Significance was determined by two-way ANOVA. $^{*}p < .05$, $^{**}p < .01$.

### Extended Data Fig. 1

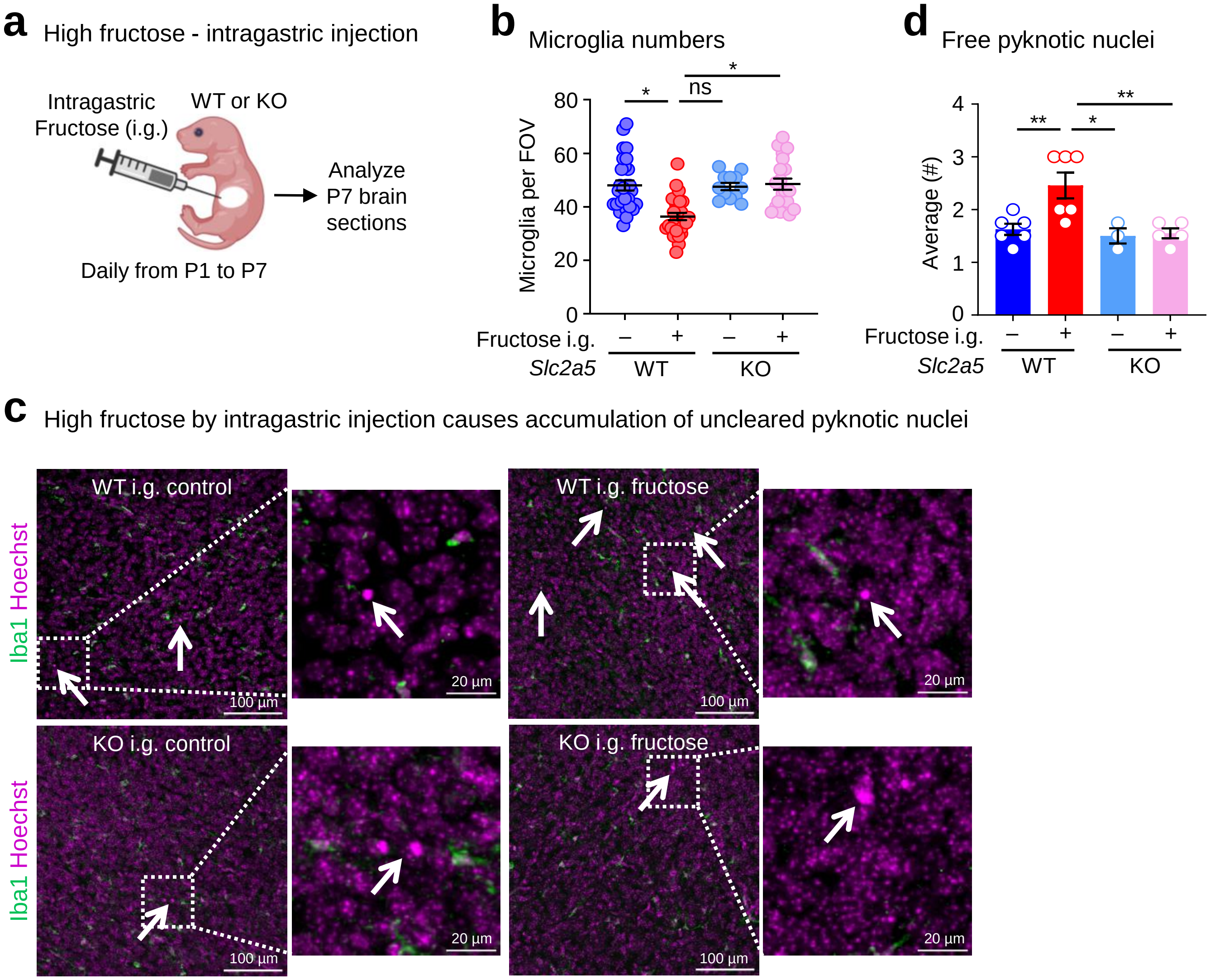

**Extended Data Fig. 2.**

**(a, b)** Schematics **(a)** and quantitation **(b)** of microglial morphological features analyzed, highlighted in red. See also **Fig. 1d** for quantification of 1-3. **(b)** Quantification of 4 and 5 from **(a)** of microglia from wildtype (WT) and *Slc2a5*$^{-/-}$ (KO) P7 neonates born to dams on control diet (CD) or high fructose diet (HF) as in **Fig. 1b**, based on four fields of view (FOVs) analyzed per mouse. Images are representative of six WT mice on CD (blue), nine WT mice on HF (red), eight KO mice on CD (light blue), and five KO mice on HF (pink). Data are shown as mean ± SEM. Significance was determined by two-way ANOVA. ns = not significant.

**Extended Data Fig. 2**

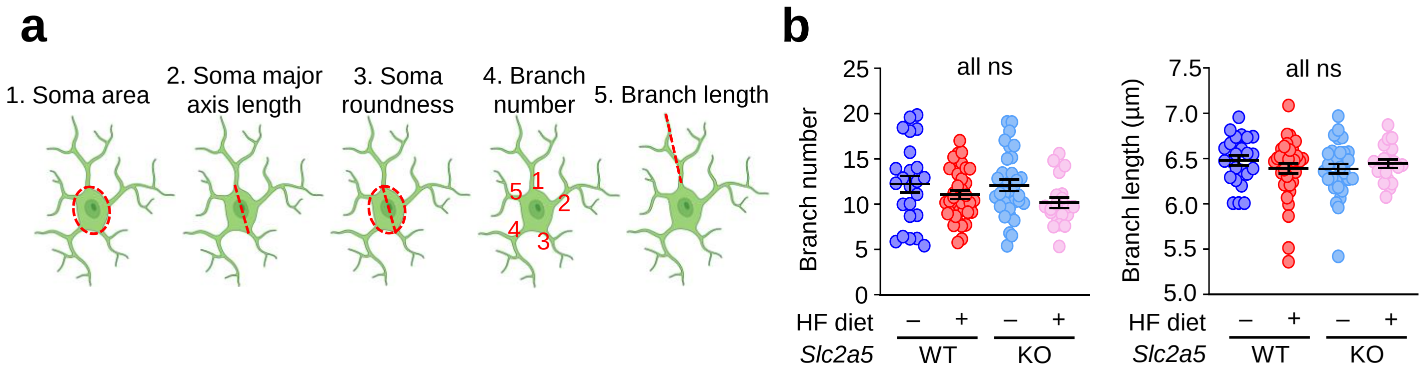

**Extended Data Fig. 3.**

**(a, b)** Representative images **(a)** and quantitation **(b)** of the cortex from wildtype (WT) and *Slc2a5*$^{-/-}$ (KO) P7 neonates exposed to maternal CD or HF in **Fig. 1b** showing Iba1 staining of microglia and Hoechst staining of nuclei. Arrows denote condensed chromatin of apoptotic cells by pyknotic nuclei accumulation. Scale bars, 100 µm (inlay, 20 µm). **(b)** Quantitation of average number of free pyknotic nuclei in **(a)**, from four FOVs per mouse with the average of those FOVs plotted per mouse. Images are representative of five WT mice on CD (blue), eight WT mice on HF (red), ten KO mice on CD (light blue), and eight KO mice on HF (pink). Data are shown as mean ± SEM. Significance was determined by two-way ANOVA. *$p < .05$, **$p < .01$, ***$p < .001$.

**(c)** Representative images of both Iba1+ microglia associated with TUNEL (bound or internalized) and free TUNEL+ puncta from experiments performed in **Fig. 1f**.

**(d)** Quantitation of the fraction of microglia containing two or more PSD95+ puncta in the prefrontal cortex from WT and KO P7 mice exposed to maternal CD or HF in **Fig. 1b**. The number of microglia with two or more PSD95+ puncta was tabulated from four FOVs per mouse for each condition and the average of those FOVs for each mouse is shown. Scale bar, 50 µm. ns = not significant, *$p < .05$, **$p < .01$.

**(e)** Mouse microglia are the only immune cell, including the only macrophage or monocyte subset, that expresses *Slc2a5*. Analysis of RNA sequencing data from ImmGen (ImmGen, 2020) showing exclusive expression of *Slc2a5* in mouse microglia across all immune cell subsets (left) and macrophage/monocyte subsets (right). Shown is data from all available immune cell and macrophage subsets.

**(f)** Microglia uniquely express *Slc2a5* or *SLC2A5* in mouse and human brains, respectively. Analysis of RNA sequencing data from Zhang, et al., 2014 and Zhang, et al., 2016 for *Slc2a5* expression in central nervous system (CNS) cells in mice and *SLC2A5* expression in CNS cells in humans. Both mouse and human microglia uniquely express *Slc2a5* and *SLC2A5* in mouse and human CNS, respectively.

Extended Data Fig. 3

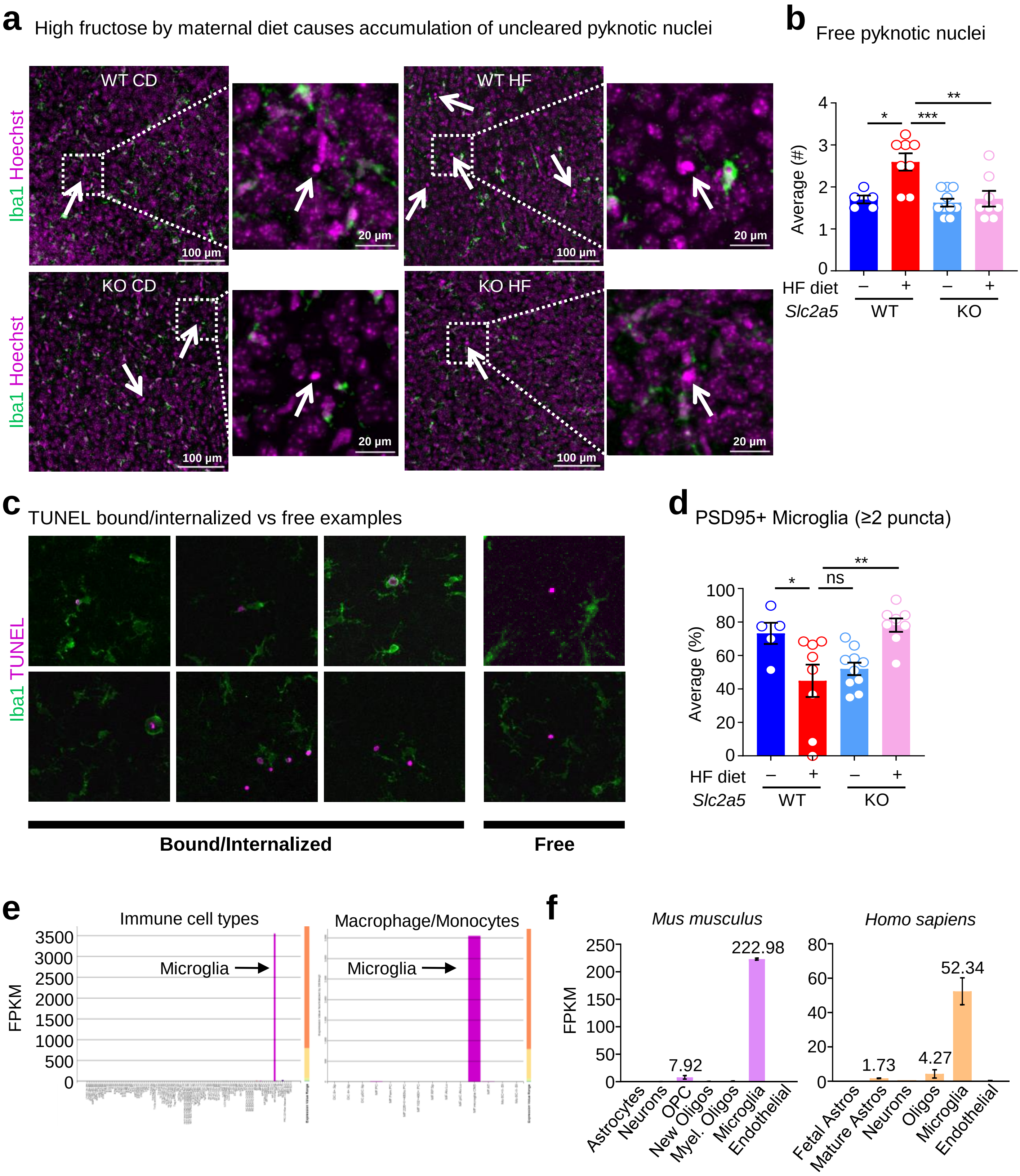

**Extended Data Fig. 4.**

**(a)** Absence of *Slc2a5* in murine microglia confirmed by RT-qPCR. All commercially available TaqMan probes were tested on P2-P4 neonatal microglia isolated from WT and *Slc2a5*$^{-/-}$ mice. Data are shown as mean ± SEM. Significance was determined using independent samples t-test. ****$p < .0001$.

**(b)** Probe coverage of murine *Slc2a5* TaqMan spanning all 14 exons. The first four exons of *Slc2a5* are deleted in *Slc2a5*$^{-/-}$ mice. Indicated probes (*) were further validated in **Fig. 2a**.

### Extended Data Fig. 4

**a** Successful deletion of *Slc2a5* from mouse microglia as shown by multiple RT-qPCR probes

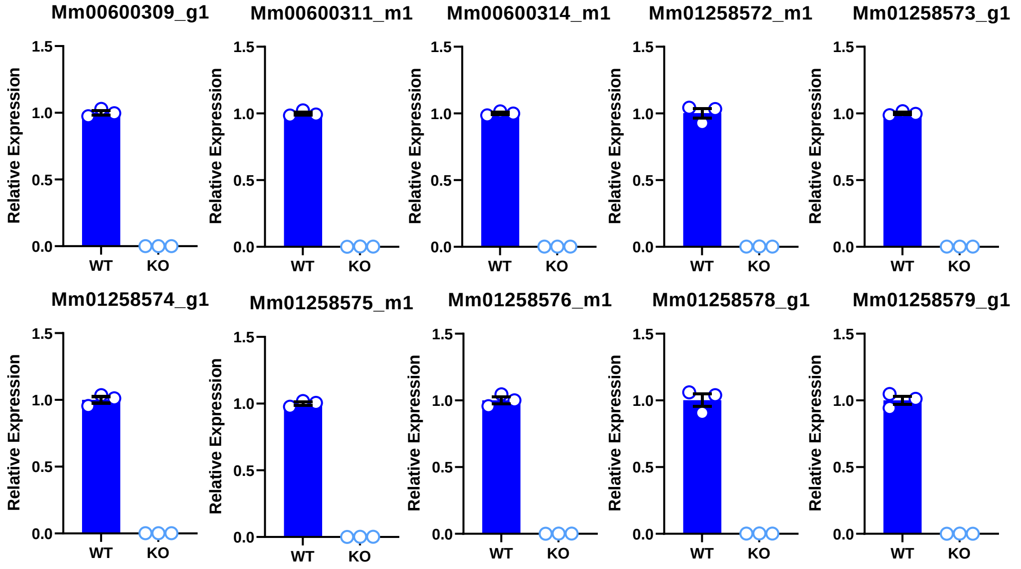


**b** Exon boundaries covered by indicated RT-qPCR probes (in red)

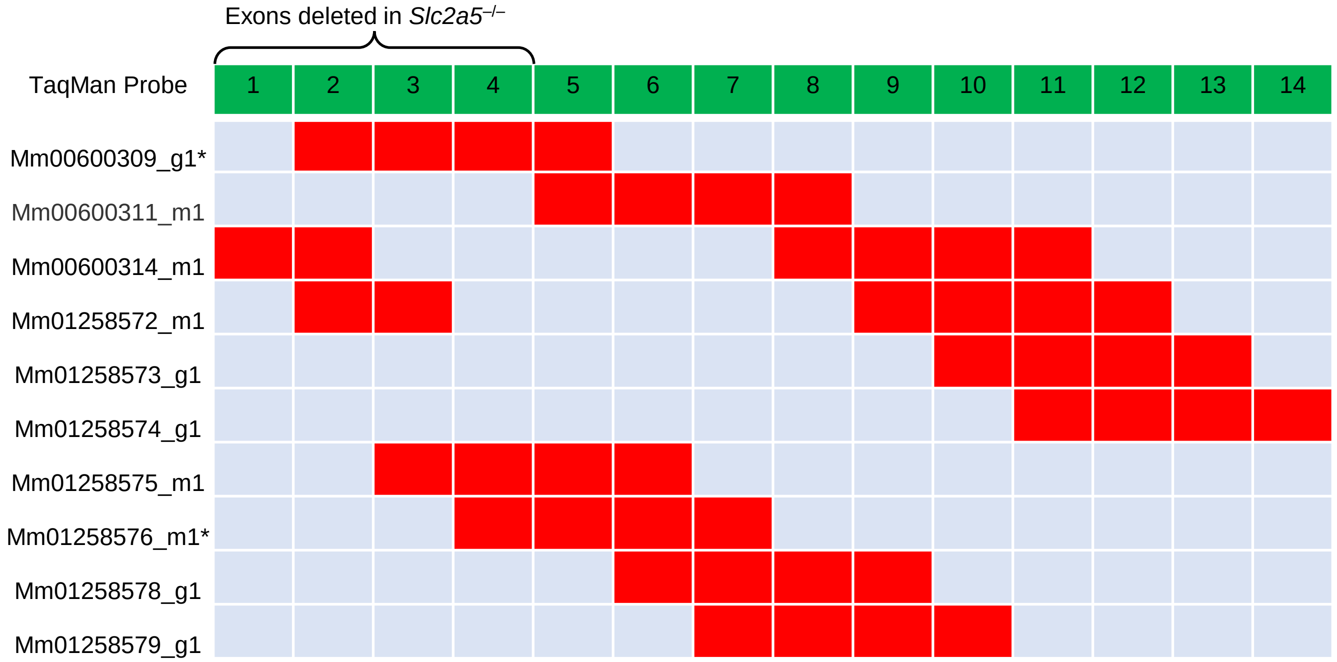

**Extended Data Fig. 5.**

**(a)** Schematic of assays performed to assess microglia phagocytosis in Fig. 2c-h. Microglia were isolated from P2-P4 WT and KO mice and cultured at physiological oxygen (3.5%) in serum-free media containing 5 mM glucose with either 0 mM fructose and 20 mM mannitol, 1 mM fructose and 19 mM mannitol, or 5mM fructose and 15 mM mannitol for one week. Synaptosomes or apoptotic neurons were labeled with CypHer5E and then cultured with conditioned microglia for 30 min. Phagocytosis was subsequently analyzed via flow cytometry or confocal microscopy.

**(b)** High fructose inhibits primary microglial expansion in mixed glial cultures. Mixed glial cultures were generated using dissected cortices from WT P2-P4 neonates. Tissue mixtures were grown in 20 mM glucose in the presence or absence of 5 mM fructose. Microglia yield was quantified after 18 d of culture. Data are from three independent experiments, shown as mean ± SEM. Significance was determined using independent samples t-test. ****$p < .0001$.

**(c)** High fructose impairs mouse primary microglia efferocytosis. Apoptotic neurons were generated by treating the neuron cell line N2A with staurosporine for 12 h, then incubated with primary microglia at a 1:1 phagocyte:target ratio for 30 min. Microglia were collected and analyzed via flow cytometry. Data is presented as phagocytic index (percent phagocytosis in experimental microglia divided by percent phagocytosis in control microglia) and is from three independent experiments with 3 or more technical replicates per condition. Data are shown as mean ± SEM. Significance was determined using independent samples t-test. ***p < .001.

**(d)** High fructose exposure enhances *Slc2a5* expression in human pluripotent stem cell (PSC)-derived microglia. Human PSC-derived microglia were cultured in complete RPMI containing 10 mM glucose with either 0 mM fructose and 20 mM mannitol, 5 mM fructose and 15 mM mannitol, or 15 mM fructose and 5 mM mannitol for 6 d and were then harvested for analysis of *Slc2a5* expression via RT-qPCR. Data are from three independent experiments with expression normalized to 18s. Data are shown as mean ± SEM. Significance was determined by one-way ANOVA. ns = not significant, ****$p < .0001$.

**(e)** High fructose impedes phagocytosis of apoptotic neurons by human PSC-derived microglia. Experiments were performed similar to **(c)** with conditions described in **(d)**, but with hPSC-derived microglia. Data are from three independent experiments with at least 3 technical replicates per condition. Data are shown as mean ± SEM. Significance was determined using independent samples t-test. ***$p < .001$.

**(f)** *Slc2a5* deletion rescues impaired phagocytosis of apoptotic neurons by primary microglia. Experiments were performed similar to **(c)**, but using primary microglia isolated from *Slc2a5*$^{-/-}$ mice. Data are from three independent experiments with at least 3 technical replicates per condition. Data are shown as mean ± SEM. Significance was determined using independent samples t-test. ns = not significant.

### Extended Data Fig. 5

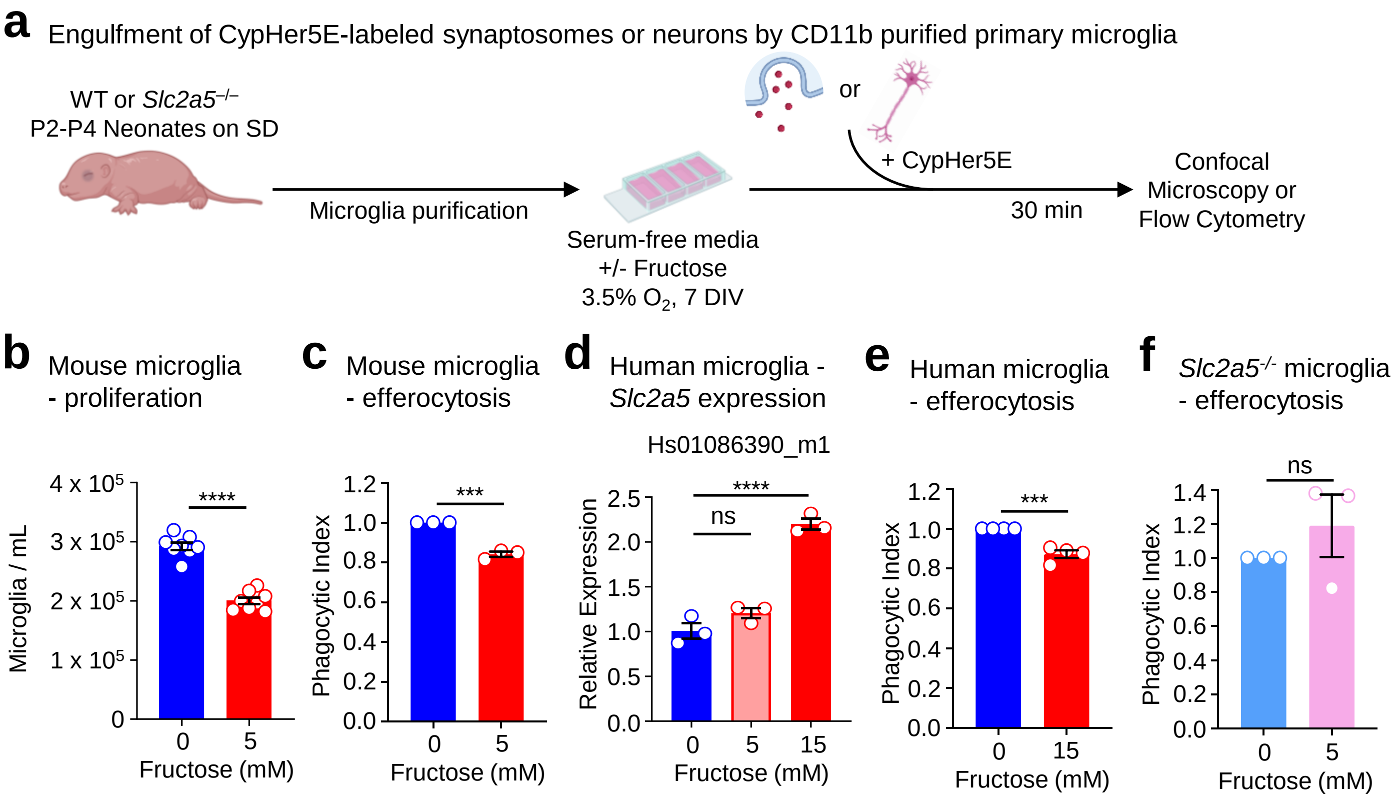

**Extended Data Fig. 6.**

**(a-c)** Relative intensity of indicated metabolites from experiments detailed in **Fig. 3f**. Data are shown as mean ± SEM. Significance was determined using two-way ANOVA. *$p < .05$, **$p < .01$, ****$p < .0001$. Fructose 6-phosphate (F6P), glyceraldehyde 3-phosphate (G3P), 3-phosphoglyceric acid (3-PG), phosphoenolpyruvic acid (PEP), nicotinamide adenine dinucleotide (NAD+), D-fructose, Lactic acid, Glutamic acid, Glutathione, and Oxidized Glutathione are shown.

### Extended Data Fig. 6

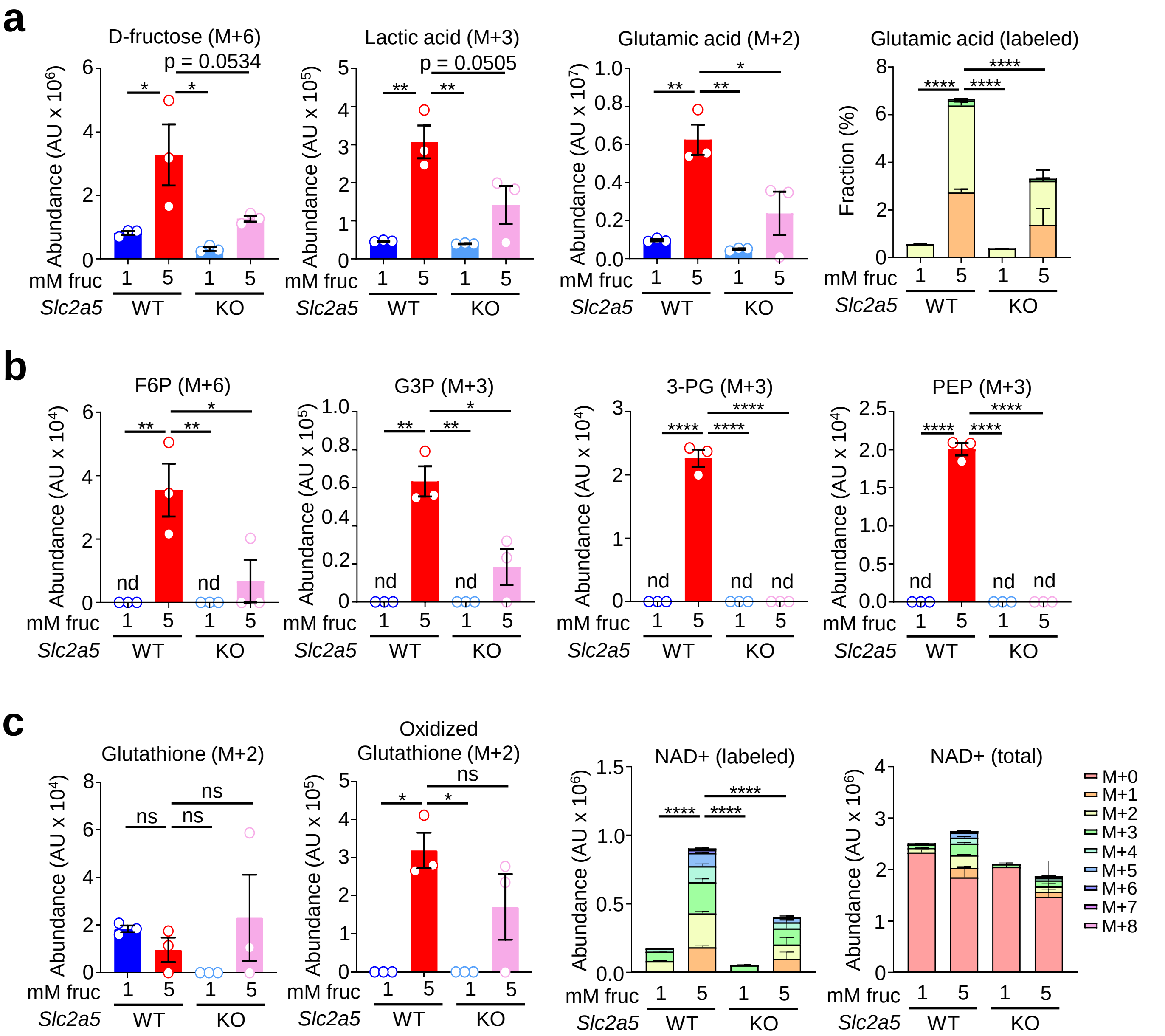

**Extended Data Fig. 7.**

Relative intensity of alanine and TCA cycle intermediates downstream of fructolysis from experiments detailed in **Fig. 3f**. Data are shown as mean ± SEM. Significance was determined using two-way ANOVA. ns = not significant, *$p < .05$, **$p < .01$, ***p < .001, ****$p < .0001$.

**Extended Data Fig. 7**

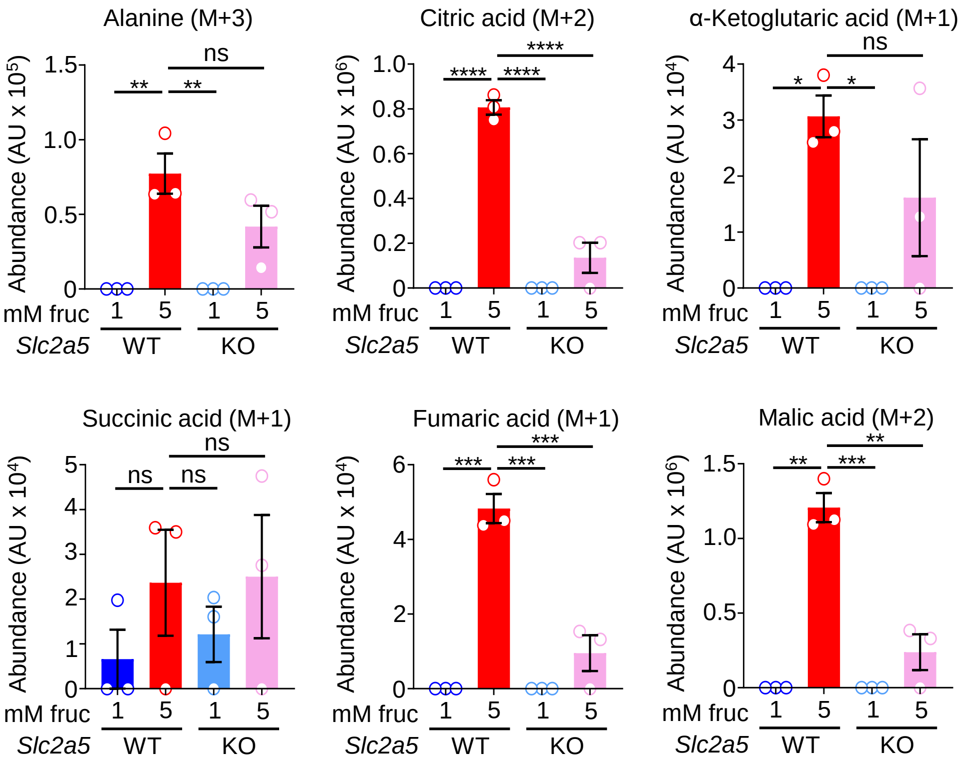

**Extended Data Fig. 8.**

Additional Modified Barnes maze performance results from experiments performed in **Fig. 4c,d**.

Data are shown as mean ± SEM. Significance was determined using two-way ANOVA. ns = not significant.

**Extended Data Fig. 8**

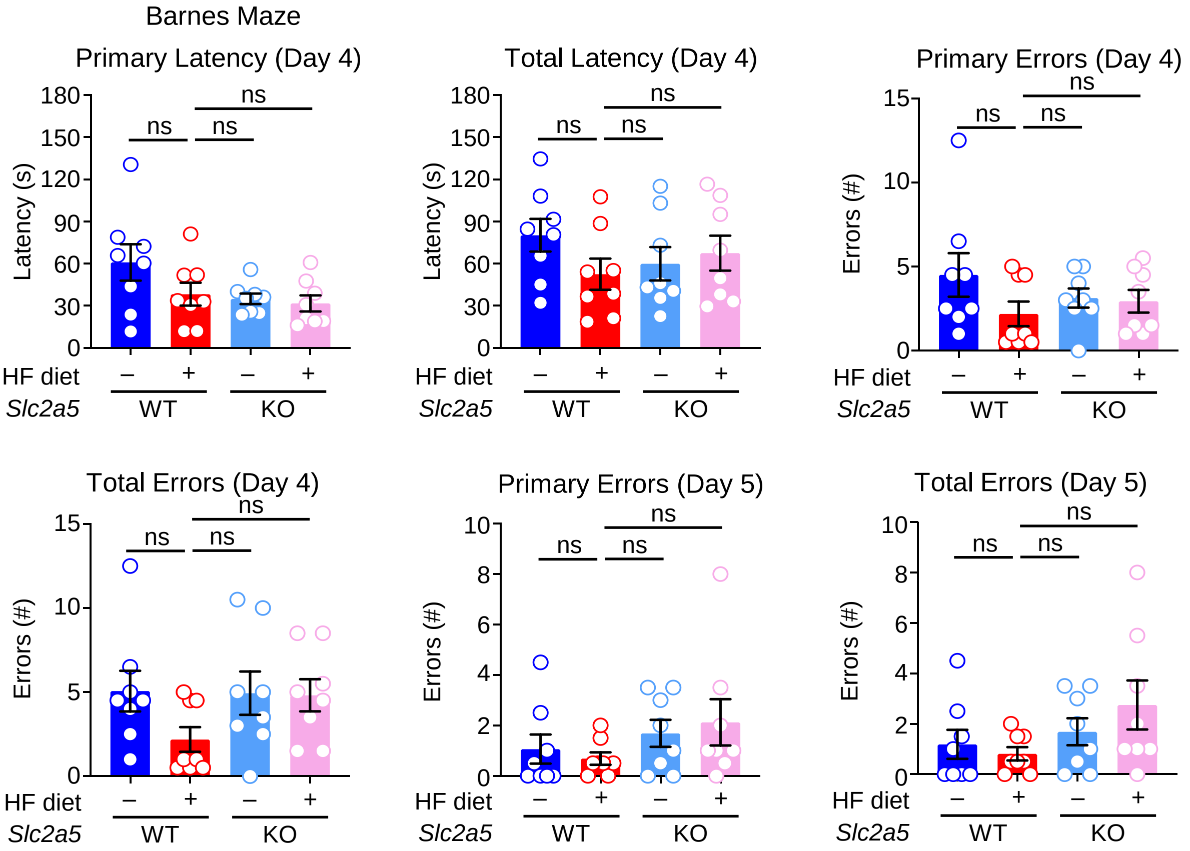
